# Mechanobiological Conditioning of Mesenchymal Stem Cells Enhances Therapeutic Angiogenesis by Inducing a Hybrid Pericyte-Endothelial Phenotype

**DOI:** 10.1101/487710

**Authors:** Jason Lee, Kayla Henderson, Miguel Armenta-Ochoa, Austin Veith, Pablo Maceda, Eun Yoon, Lara Samarneh, Mitchell Wong, Andrew K. Dunn, Aaron B. Baker

## Abstract

Stem cell therapies have great promise for revolutionizing treatments for cardiovascular disease and other disorders but have not yet achieved their potential due to poor efficacy and heterogeneity in patient response. Here, we used a novel high throughput screening system to optimize the conditioning of mesenchymal stem cells using a combinatorial set of biochemical factors, pharmacological inhibitors and biomechanical forces. Our studies revealed that a combination of specific kinase inhibitors and a complex mechanical strain waveform dramatically increased the population of mesenchymal stem cells that express markers for both pericytes and endothelial cells. These mechanically and pharmacologically conditioned mesenchymal stem cells had superior properties in enhancing endothelial tube formation, production of angiogenic growth factors and induction of angiogenesis following implantation. Overall, our work supports that combinatorial optimization of mechanical conditioning and pharmacological treatments can significantly enhance the regenerative properties of mesenchymal stem cells.

---

Cell based therapies have great potential for revolutionizing the treatment of diseases that are not amenable to traditional treatments. Therapies based on mesenchymal stem cells (MSCs) are particularly appealing as they are a source of autologous cells with diverse multipotency and can be harvested from patients with relative ease. In addition, MSCs are able to self-renew and have immunosuppressive properties that make them ideal candidates for autologous cellular therapeutics^1^. For cardiovascular therapies, MSCs have been explored for the treatment of myocardial infarct and peripheral ischemia^2–7^. However, these trials have not shown consistent long-term benefits to patients from MSC therapies in spite of intense investigation by many groups^8–11^. MSCs have several aspects that limit their potential for use in cell therapies. In conventional culture conditions, MSCs lose their differentiation potential and have reduced therapeutic properties after expansion^12,13^. In addition, the isolated MSC populations have a high degree of heterogeneity, with subsets of cells that have varying degrees of potential for inducing regeneration^14,15^. Moreover, the therapeutic potential of MSCs is altered by the health of the patient from which they are harvested. This is a major limitation as patients with advanced age, obesity, diabetes and other chronic disorders have MSCs with altered differentiation potential and regenerative properties^12,13,16–19^. Thus, those patients who would likely benefit the most from cell therapies are those who have the least regenerative MSCs. Genetic modification of MSCs could address some of these issues but also raise concerns of tumorigenicity^20^. Consequently, there is an intense interest in identifying alternative strategies for making MSCs more effective and reliable in spite of patient-to-patient differences in MSC behavior or reduced regenerative capability.

The differentiation of MSCs into endothelial cells would be highly advantageous for many clinical applications including the treatment of ischemia and endothelialization of vascular grafts. Unfortunately, there has not been consensus in the field as to what conditions induce endothelial differentiation in MSCs or even if the phenotype obtained is truly endothelial in nature^21–24^. Biophysical forces, including shear stress and mechanical stretch, have been used to condition MSCs into vascular phenotypes^25–30^. Several studies have shown that shear stress and treatments such as VEGF can lead to the expression of endothelial markers in MSCs^21,23^. However, some studies have found contrasting results in endothelial differentiation for identical treatments or an increase in markers for vascular smooth muscle cells (vSMCs) in addition to those for endothelial cells^29–32^. Thus, it is unclear what conditions are optimal for obtaining an endothelial cell-like phenotype in MSCs. Moreover, it is unclear if enhancing endothelial or vascular phenotype also increases the effectiveness of MSC therapies.

In this work, we used a novel high throughput system for applying mechanical stretch to cultured cells to explore the ability of applied mechanical forces to enhance the regenerative properties of MSCs. Using this system, we performed a set of powerful screening assays that explored the synergy between biochemical factors, pharmacological inhibitors and biomechanical forces in conditioning MSCs into vascular cell-like phenotypes that have enhanced regenerative properties. We identified specific mechanical conditions that induce maximal activation of the Hippo and Smad signaling pathways in MSCs. Using a high throughput screen incorporating applied mechanical load, we also identified several pharmacological inhibitors that synergistically activated these pathways. Surprisingly, a rigorous analysis of the phenotype of the cells treated with combined mechanical conditioning and drug treatments revealed increased expression both pericyte and endothelial markers. These optimally conditioned MSCs exhibited enhanced pericyte and endothelial behavior ***in vitro,*** production of angiogenic soluble factors, and increased therapeutic potential in enhancing blood vessel growth in animal models of subcutaneous implantation and hind limb ischemia. Taken together, our study demonstrates that the regenerative potential of MSCs can be markedly improved by optimized mechanical and pharmocological conditioning. Further, we identify that MSCs can be stimulated with complex mechanical loads to adopt a hybrid phenotype that has combined properties of pericytes/endothelial cells and increased regenerative capacity.

## Results

### Mechanical regulation of transcription factor activity and signaling pathways in mesenchymal stem cells

We explored whether there was synergy between biochemical and mechanical stimuli to activate the transcription factors by exposing the cells to VEGF-A or TGF-β1 during mechanical loading. These two factors have been linked to stimulating differentiation in MSCs to vascular cell types and other lineages^21,23,33^. We transduced MSCs with lentiviruses expressing luciferase reporter constructs for transcription factors including FOXO, TCF/LEF, Smad2/3 and SRE and selected these to obtain a stable reporter cell line. All mechanical loading studies were performed using a high throughput biaxial oscillatory strain system (HT-BOSS) recently developed by our group.^34^ This system allows the simultaneous application of mechanical forces to up to 576 wells of cultured cells in a standard 96 well format. For the transcription factor FOXO, we observed no stimulation of transcription factor activity, but instead found a reduction in FOXO activity with VEGF treatment at 2 Hz frequency of loading (5% maximal strain) or with VEGF treatment at 1 Hz frequency of loading (**Fig. 1A**). In contrast, TCF/LEF was synergistically activated by loading at 0.1 Hz in combination with VEGF treatment (**Fig. 1A**). The Smad2/3 transcription factors were also synergistically activated by TGF-β1 treatment and loading at 0.1 Hz, and suppressed by loading at 1 or 2 Hz (**Fig. 1A**).

**Figure 1.**
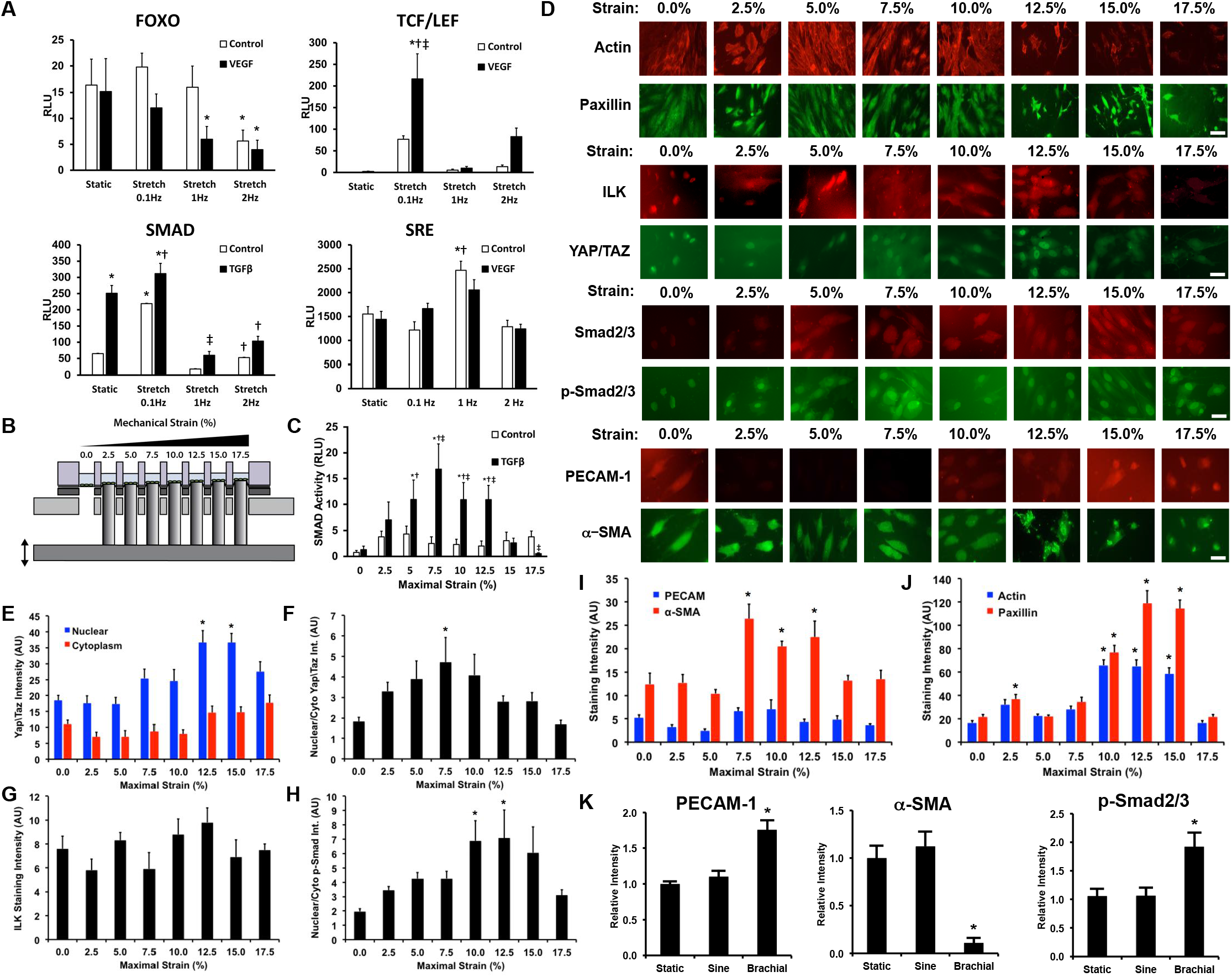
Mechanotransduction in MSCs is dependent on the magnitude and frequency of mechanical loading. (A) Transcription factor activity in MSCs was measured using a luciferase reporter assay after the application of cyclic mechanical strain for 8 hours with co-treatment with 10 ng/ml VEGF-A or 10 ng/ml TGF-β1. *\*p* < 0.05 versus static control group. † *p* < 0.05 versus static growth factor treated group. ‡*p* < 0.05 versus control group under the same mechanical loading conditions. (B) The height of the pistons is changed to apply varying mechanical strain across the plate. (C) Smad transcription factor activity in MSCs with application of load for 24 hours using the multi-strain configuration. *\*p* < 0.05 versus static control group. †*p* < 0.05 versus static growth factor treated group. ‡*p* < 0.05 versus control group under the same mechanical loading conditions. (D) The MSCs were treated with mechanical load using the multi-strain format at 0.1 Hz for 24 hours and then immunostained for markers of vascular cell differentiation or signaling pathway activation. Scale bar = 100 μm. (E) Quantitative analysis of Yap/Taz nuclear localization after mechanical loading for 24 hours. *\*p* < 0.05 versus static group. (F) Ratio of nuclear to cytoplasmic Yap/Taz in the mechanically loaded MSCs. *\*p* < 0.05 versus static group. (G) Quantification of immunostaining for ILK in MSCs treated with mechanical load for 24 hours. (H) Ratio of nuclear to cytoplasmic p-Smad2/3 in MSC after mechanical loading for 24 hours. (I) Quantification of vascular marker expression in MSCs following 24 hours of mechanical loading. *\*p* < 0.05 versus static group. (J) Quantification of focal adhesion and cytoskeletal components following mechanical loading for 24 hours. *\*p* < 0.05 versus static group. (K) Quantification of western blotting for vascular markers and signal activation. *\*p* < 0.05 versus static and sine groups.

### Multi-strain mechanical loading assays reveal optimal conditions for Smad2/3 transcription factor activation and regulation of vascular cell markers in MSCs

To optimize the magnitude of mechanical strain to maximize activation of Smad2/3, we adjusted the heights of the individual pistons to simultaneously apply a range of mechanical strains from 0 to 17.5% (**Fig. 1B**). We applied mechanical strain over this range at a frequency of 0.1 Hz and found optimal synergistic activation of Smad transcription factors at 7.5% strain with TGF-β1 co-treatment (**Fig. 1C**). We next examined the regulation of differentiation-related pathways in MSCs by loading the cells for 24 hours under multi-strain conditions at 0.1 Hz and performed immunostaining. We found a maximal increase in nuclear to cytoplasmic Yap/Taz ratio at 7.5% strain and maximal nuclear localization of Smad2/3 at 10-12.5% strain (**Fig. 1D-F**). We co-stained for integrin-linked kinase (ILK), a known suppressor of the Hippo pathway,^35^ but did not see a significant difference with mechanical load (**Fig. 1G**). We found a dose dependent increase in nuclear phospho-Smad2/3 (p-Smad2/3), with a maximal response at 10 or 12.5% strain (**Fig. 1H**). In addition, there was a maximal increase in α-SMA expression in the range of strains from 7.5-12.5% strain (**Fig. 1F, H**). The HT-BOSS system uses a linear motor that allows loading with complex physiological waveforms that simulate the loads in the body. We used this capability to reproduce the complex strain waveform that occurs in the brachial artery of the body^34,36^. We repeated the mechanical loading at 7.5% strain with 0.1 Hz frequency using both sine and brachial waveforms, and then performed western blotting. With brachial waveform loading, we found increases in PECAM-1 and p-Smad2/3 with a reduction in α-SMA expression (**Fig. 1I, J; Supplemental Fig. 1**).

### High throughput drug screening assay with mechanical load identifies kinase inhibitors that enhance Smad2/3 signaling and Hippo pathway activation in synergy with mechanical load

We next used the HT-BOSS system to perform a compound screening assay to identify compounds that could enhance Smad and Hippo pathway activation in combination with mechanical load. We performed mechanical loading on MSCs in the 96-well format in the presence of one of 80 compounds from a kinase inhibitor library. After 24 hours, we immunostained for p-Smad2/3 and Yap/Taz, and then quantified the nuclear staining of both signaling intermediates (**Fig. 2A, B; Supplemental Fig. 2, 3**). From this assay, we identified compounds that markedly increased both nuclear Yap and p-Smad2/3 (**Fig. 2**). Many of the top hits from the assay included inhibitors of the EGFR pathways (**Fig. 2C**). We chose the EGFR/ErbB-2/4 inhibitor N-(4-((3-chloro-4-fluorophenyl)amino)pyrido[3,4-d]pyrimidin-6-yl)2-butynamide (CAS 881001-19-0) and the PKCbII/EGFR inhibitor 4,5-bis(4-fluoroanilino)-phthalimide (CAS 145915-60-2) for further study based on the maximal activation of both pathways or maximal activation of the Hippo pathway, respectively.

**Figure 2.**
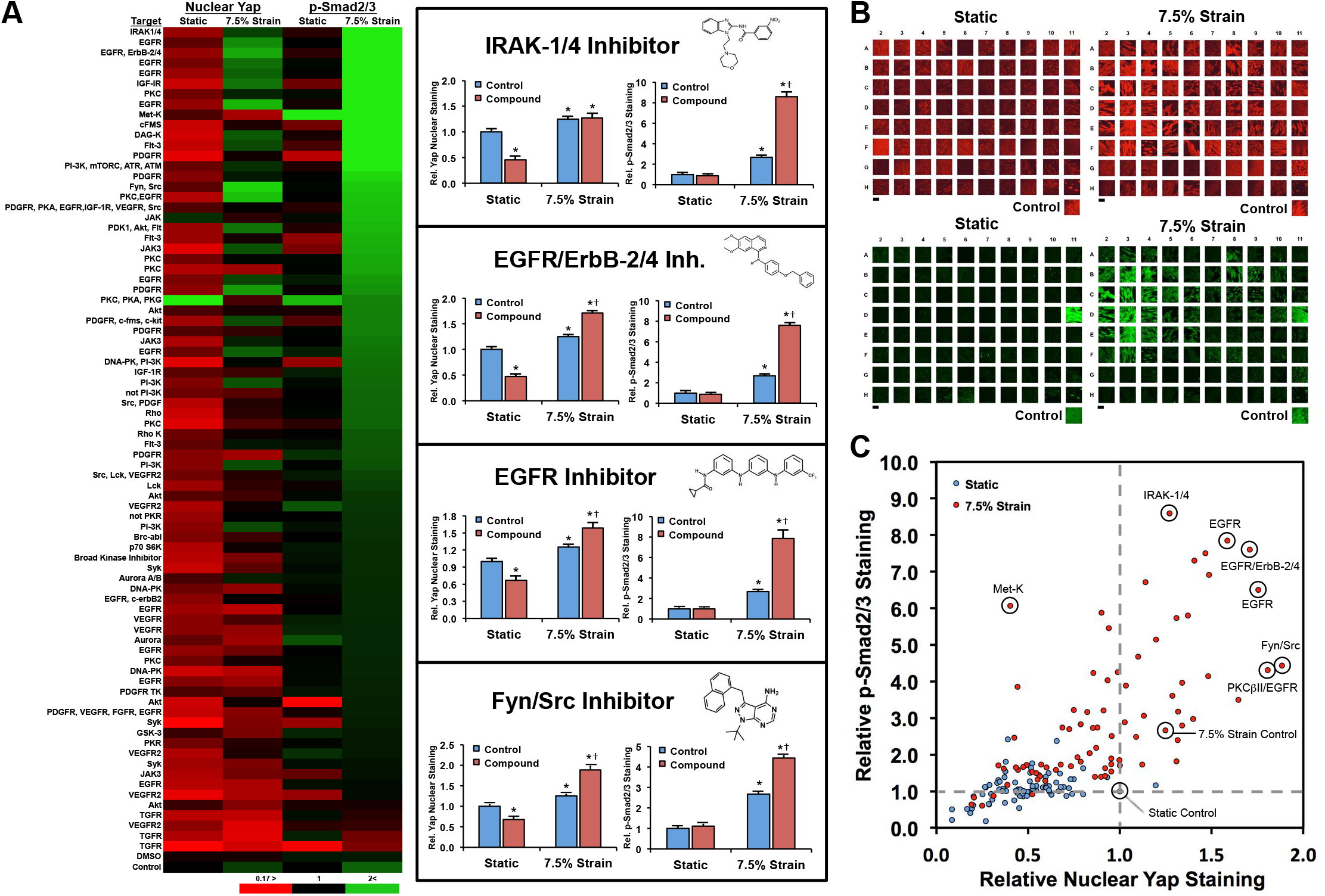
High throughput mechanobiological screen for small molecule inhibitors that have synergistic activation of Yap/Taz and Smad2/3 with mechanical loading. The MSCs were treated with 7.5% mechanical strain at 0.1 Hz for 24 hours in the presence of compounds from a library of kinase inhibitors. (A) The cells were immunostained and quantified for nuclear localization of Yap/Taz and p-Smad2/3. \**p* < 0.05 versus cells treated with static conditions under control treatment. †*p* < 0.05 versus mechanically strained control group. (B) Images from immunostaining of cells arranged in the 96 well plate format. Scale bar = 50 μm (C) Overall summary of the mechanobiological screen separated to show the response distribution for the compounds. Labeled samples indicate the target of the inhibitor used or control treatments.

### Long term treatment with mechanical load and differentiation factors regulates pluripotency markers, vascular cell markers and signaling pathways in MSCs

To examine the ability of long term application of mechanical loading to condition MSCs, we exposed the cells with treatments known to cause differentiation into vascular/cardiovascular phenotypes including 5-Azacitidine (5-Aza), dexamethasone (Dex) with insulin (Ins), PDGF-BB, hydrocortisone (HC), EGF, VEGF-A and TGF-β1^37^. After seven days of loading at 0.1 Hz with 7.5% strain for 4 hours per day, we found increased polymerized actin after loading in control media and higher levels of polymerized actin with all of the differentiation treatments under static conditions except TGF-β1 (**Supplemental Fig. 4**). Similarly, there were increased paxillin levels for the treatments 5-Aza, Dex + Ins, EGF and VEGF-A with and without load (**Supplemental Fig. 4**). Paxillin was not increased in static or loaded conditions with TGF-β1 treatment. In addition, we found that mechanical loading increased Smad2/3 and p-Smad2/3 and nuclear Yap/Taz under all the treatments (**Fig. 3A, B**). Nuclear localization of Yap/Taz was increased with mechanical loading under the majority of the treatments, with only TGF-β1 treatment markedly reducing this effect.

**Figure 3.**
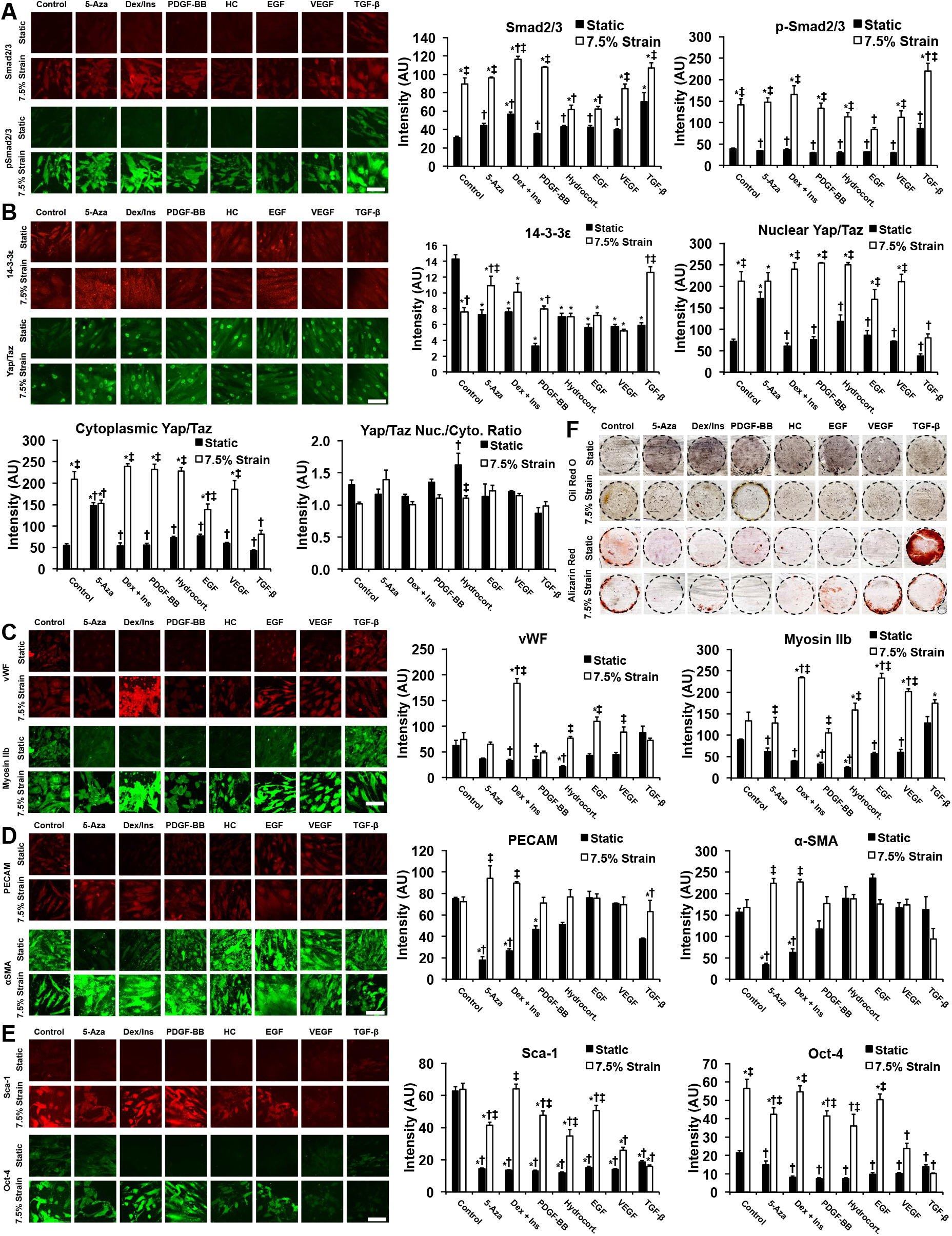
Activation of mechanotransduction pathways and expression of vascular cell markers in mesenchymal stem cells under long-term treatment with biochemical differentiation treatments and mechanical loading. MSCs were treated with differentiation treatments for seven days and then immunostained for markers of Hippo pathway/TGF-β signaling and differentiation into vascular phenotypes. Treatments included 5-Aza, dexamethasone and insulin (Dex + Ins), PDGF-BB, hydrocortisone (HC), EGF, VEGF-A and TGF-β1. (A) Immunostaining and quantification of activation of Smad2/3 after seven days of treatment. (B) Alterations in Yap/Taz and 14-3-3ε with mechanical loading and differentiation treatments. (C-E) Immunostaining for endothelial cell, vSMC markers and pluripotency markers after seven days of conditioning with mechanical load. *\*p* < 0.05 versus cells treated with static conditions under baseline treatment. †*p* < 0.05 versus mechanically strained group under baseline treatment. ‡*p* < 0.05 versus cells under static conditions with the same biochemical treatment. Scale bar = 100 μm. (F) Osteogenic and adipogenic differentiation markers of static and stretched MSCs were measured after the differentiation treatments through Alizarin Red S and Oil Red O staining.

We next examined the expression of markers for endothelial cells, vSMCs and pluripotency using immunostaining. Mechanical loading increased the endothelial marker Von Willebrand factor (vWF) when combined with dexamethasone and insulin treatment, and to a lesser extent with EGF and VEGF-A treatment (**Fig. 3C**). In addition, mechanical load increased PECAM-1 in combination with 5-Aza and with dexamethasone and insulin treatment (**Fig. 3D**). However, these treatments in most cases led to an increase or no reduction in markers for vSMCs, including Myosin IIb and α-SMA (**Fig. 3C, D**). Oct-4 is a marker associated with the maintenance of pluripotency in MSCs^38^. Under the treatments without mechanical load, there was a reduction in Oct-4 expression (**Fig. 3E**). However, with mechanical loading, Oct-4 was markedly increased in the mechanically loaded groups for all treatments except VEGF-A and TGF-β1. Sca-1 has been used as a marker to identify stem cells in many tissues, although the exact nature of isolated Sca-1+ cells remains controversial in terms of the stemness and pluripotency^39^. Expression of Sca-1 has been linked functionally to cardiac repair both in knockout models and in delivery of Sca-1+ cells^40,41^. Biochemical treatments alone resulted in decreased Sca-1 expression, while co-treatment in combination with mechanical load increased Sca-1 expression in all treatments except TGF-β1 (**Fig. 3E**). Staining for adipogenic and osteogenic differentiation demonstrated that there was little adipogenesis under any of the treatments (**Fig. 3F**). Osteogenic differentiation was observed with TGF-β1 treatment under static conditions but did not occur under TGF-β1 treatment with mechanical loading (**Fig. 3F**).

### Brachial waveform mechanical loading and pharmacological inhibition induce a hybrid endothelial/pericyte phenotype in MSCs with enhanced angiogenic properties

The induction of endothelial phenotype in MSCs would be advantageous in many therapeutic applications. Unfortunately, several studies of endothelial differentiation of MSCs have shown contradictory results^21,23,24^. Recent studies have also suggested that MSCs may be related to or identical to pericytes^42^. However, other studies do not support that pericytes have stem cell-like properties^43^. Pericyte markers in MSCs are correlated with enhanced regenerative properties in MSCs^44–47^. Many studies have been done that only look at one or two markers and conclude a phenotype of endothelial and vSMC lineage, making the true phenotype difficult to assess. To address these issues, we applied a combinatorial set of mechanical loading and biochemical/pharmacological treatments to MSCs and then assessed their phenotype with multiple markers using flow cytometry (**Supplemental Fig. 5**). We included in the treatments the pharmacological inhibitors identified in the high throughput screen to synergistically activate p-Smad2/3 and induce Yap/Taz nuclear localization. Using this relatively rigorous definition of endothelial phenotype, there was little endothelial lineage expressed by the cells under baseline conditions and with the treatments. Notably, VEGF did not increase the endothelial lineage in the MSCs, as defined by multiple markers using flow cytometry. With brachial waveform mechanical loading, there was an increase in a mixed cell phenotype that had both increased endothelial markers PECAM-1 and markers for pericytes including CD146, Nestin and PDGFRβ (**Fig. 4A; Supplemental Fig. 6**). This population was also largely positive for NG2, suggesting a hybrid phenotype of type 2 pericytes and endothelial cells (**Fig. 4A; Supplemental Fig. 7A**). There was a similar enrichment in the population of cells that expressed additional endothelial markers and pericyte markers (PECAM^+^ CD105^+^ VE-Cad^+^ CD146^+^ Nestin^+^ PDGFRβ^+^ NG2^+^) following treatment with brachial loading and co-treatment with kinase inhibitors identified from drug screening assay (**Supplemental Fig. 7B**). To analyze the pericyte phenotype, we measured cells that were positive for pericyte markers and negative for endothelial cells markers (PECAM^-^ CD105^-^ VE-Cad^-^ CD146^+^ Nestin^+^ PDGFRβ^+^ cells; **Supplemental Fig. 7C**), which represented only a small subset of the overall cell population.

**Figure 4.**
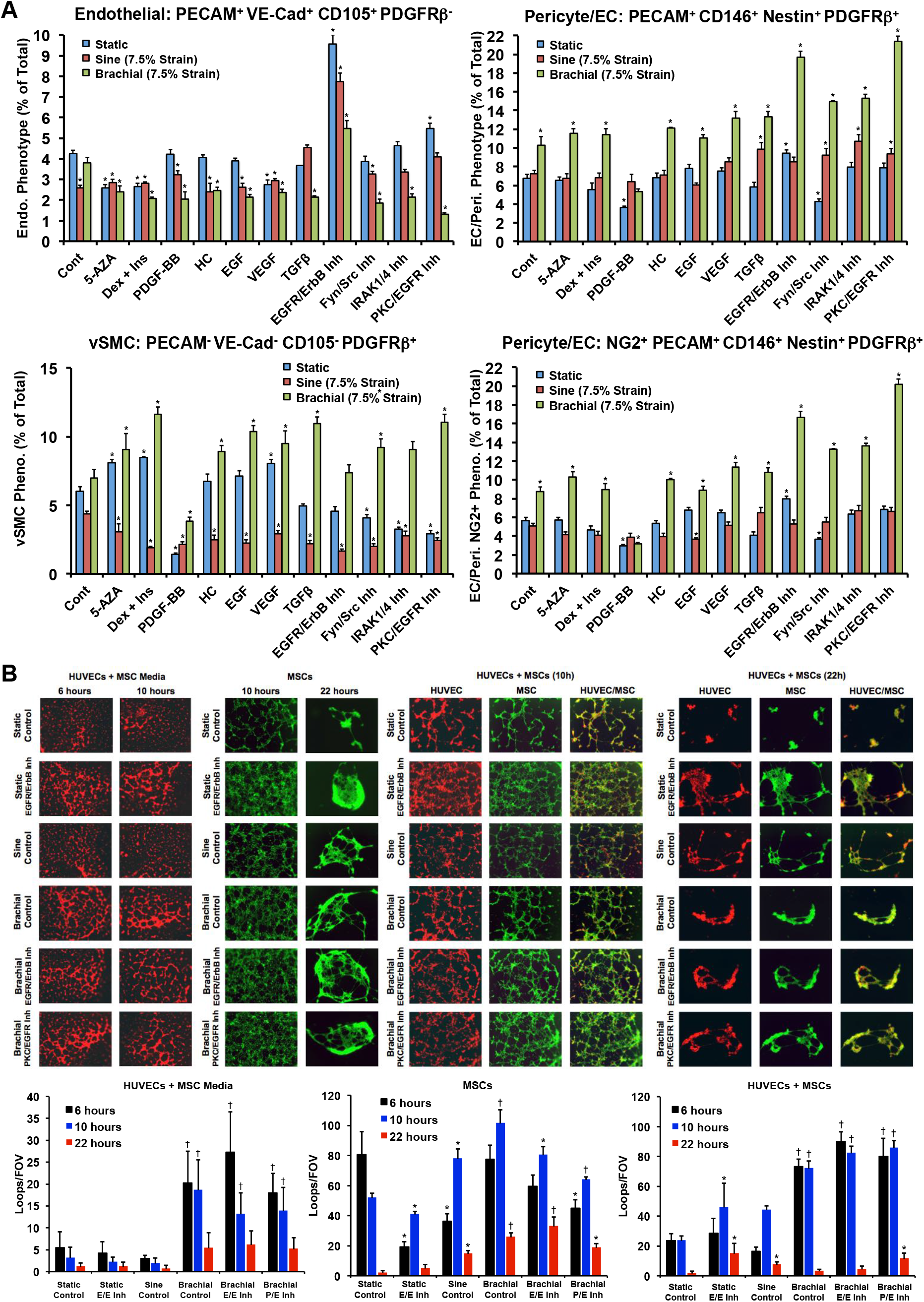
Biomechanical stimulation of mesenchymal stem cells with the brachial waveform and specific small molecule inhibitors leads to increased expression of endothelial cell and pericyte markers and enhanced pericyte-like activity. (A) Analysis of cells treated with biochemical factors and/or small molecule inhibitors for seven days with 4 hours a day of mechanical loading. Cells were labeled for multiple markers of endothelial and pericyte lineage and analyzed using flow cytometry. *\*p* < 0.05 versus control/non-loaded group. (B) Tube formation assay analyzing the activity of the conditioned media derived from MSCs under the treatments (HUVECs + MSC Media), the tube formation activity of the MSCs on Matrigel (MSCs), or the tube formation activity in MSCs seeded on Matrigel in co-culture with endothelial cells (HUVECs + MSCs). *\*p* < 0.05 versus static group at the same time point. †*p* < 0.05 versus static and sine group at the same time point. Scale bar = 100 μm.

### Biomechanical conditioning increases pericyte-like activity and angiogenic properties of

MSCs. We examined whether the cells under the different conditions had increased pericyte-like behavior and pro-angiogenic activity using a tube formation assay. Conditioned media from MSCs treated with brachial loading with or without pharmacological inhibitors induced increased tube formation in endothelial cells (**Fig. 4B**). We also plated the MSCs on Matrigel directly and found that there was increased formation of mature tubes at later time points (**Fig. 4B**). We then mixed MSCs with endothelial cells and plated them together to examine the effect of mechanical/biochemical conditioning on inducing pericyte-like behavior in MSCs. This analysis showed that MSCs exposed to brachial waveform loading had increased tube formation in co-culture with endothelial cells (**Fig. 4B**). We performed mechanical loading of MSCs with the sine and brachial waveforms (4 hours/day) and treated with pharmacological inhibitors for seven days (the compounds were not added in the final day treatment). The conditioned media from MSCs under the treatments were found to decrease endothelial cell proliferation in comparison to conditioned media from cells grown under control conditions (**Supplemental Fig. 8**).

We next examined the production of growth factors by MSCs under mechanical loading with co-treatment with pharmacological inhibitors following short term (24 hours) or longer term (7 days) treatment. We performed an antibody array analysis on the conditioned media from the treated cells and identified alterations in growth factor production by the MSCs after 24 hours or seven days of loading with co-treatments (**Supplemental Fig. 9**). We used ELISA to measure changes in the growth factors and found increases in angiotensin-1 (Ang-1) and VEGF-A with sine wave loading after 24 hours (**Supplemental Fig. 10**). Following seven days of loading, there was a marked increase in hepatocyte growth factor (HGF), a known angiogenic factor,^48^ in all of the groups treated with mechanical load (**Supplemental Fig. 11**). In the groups treated with brachial loading, there was also a decrease in TGF-β1, Ang-1 and Ang-2 (**Supplemental Fig. 11**).

### Gene expression analysis by RNA-Seq demonstrates enhancement of pericyte/endothelial phenotypes and pro-angiogenic gene expression in MSCs with optimized mechanical conditioning and small molecule treatment

To further examine whether mechanical loading/drug treatment enhanced angiogenic and pericyte-like phenotype in MSCs, we performed RNASeq on MSCs under the various treatments. A differential gene expression analysis revealed that treatment with the brachial waveform mechanical loading, with or without the ErbB/EGRB (E/E) inhibitor, significantly regulated a large number of genes in comparison to the sine wave form (1,010 or 878 genes for brachial or brachial + E/E inhibitor, respectively, versus 130 genes for sine loading treated cells; **Fig. 5A**). Treatment with the E/E inhibitor also modulated a relatively large number of genes (723 genes; **Fig. 5A; Supplemental Fig. 12**). Cells treated with brachial loading or brachial loading with E/E inhibitor treatment had similar patterns of gene expression while MSCs treated with sine or static conditions were more similar (**Fig. 5B**). To test whether the MSCs under the treatment were developing a more pericyte-like phenotype, we compared the gene expression of the cells in our study to the results of a gene expression analysis from another study that examined the development of pericytes and vSMCs from mesenchymal cells.^49^ For brachial and brachial + E/E treated cells, there were gene expression changes consistent with a type 1 pericyte phenotype (**Fig. 5C, D**). Gene ontology analysis revealed that the majority of the most significantly upregulated gene ontology groups were related to angiogenesis for both the brachial and brachial with E/E inhibitor groups but not for other groups (**Fig. 5E**). We examined a set of genes associated with the development of endothelial phenotype^50^ and found that some but not all of these genes were strongly upregulated (**Fig. 5F; Supplemental Fig. 13**). We also examined the gene expression for soluble factors related to angiogenesis and found increases in ANGPT2 and HGF for static + E/E inhibitor, brachial and brachial + E/E inhibitor-treated cells (**Supplemental Fig. 14**). To examine whether there were other types of differentiation occurring, we examined gene expression relating to osteogenesis,^51^ adipogenesis,^51^ and chondrogenesis.^52^ There was no broad increase in adipogenic or chondrogenic phenotypes (**Supplemental Fig. 15 and 16**). There were significant increases in some of the osteogenesis genes for the brachial and brachial + E/E inhibitor groups (**Supplemental Fig. 17**). However, many of these genes also have expression/roles in endothelial behavior or in ECM remodeling.

**Figure 5.**
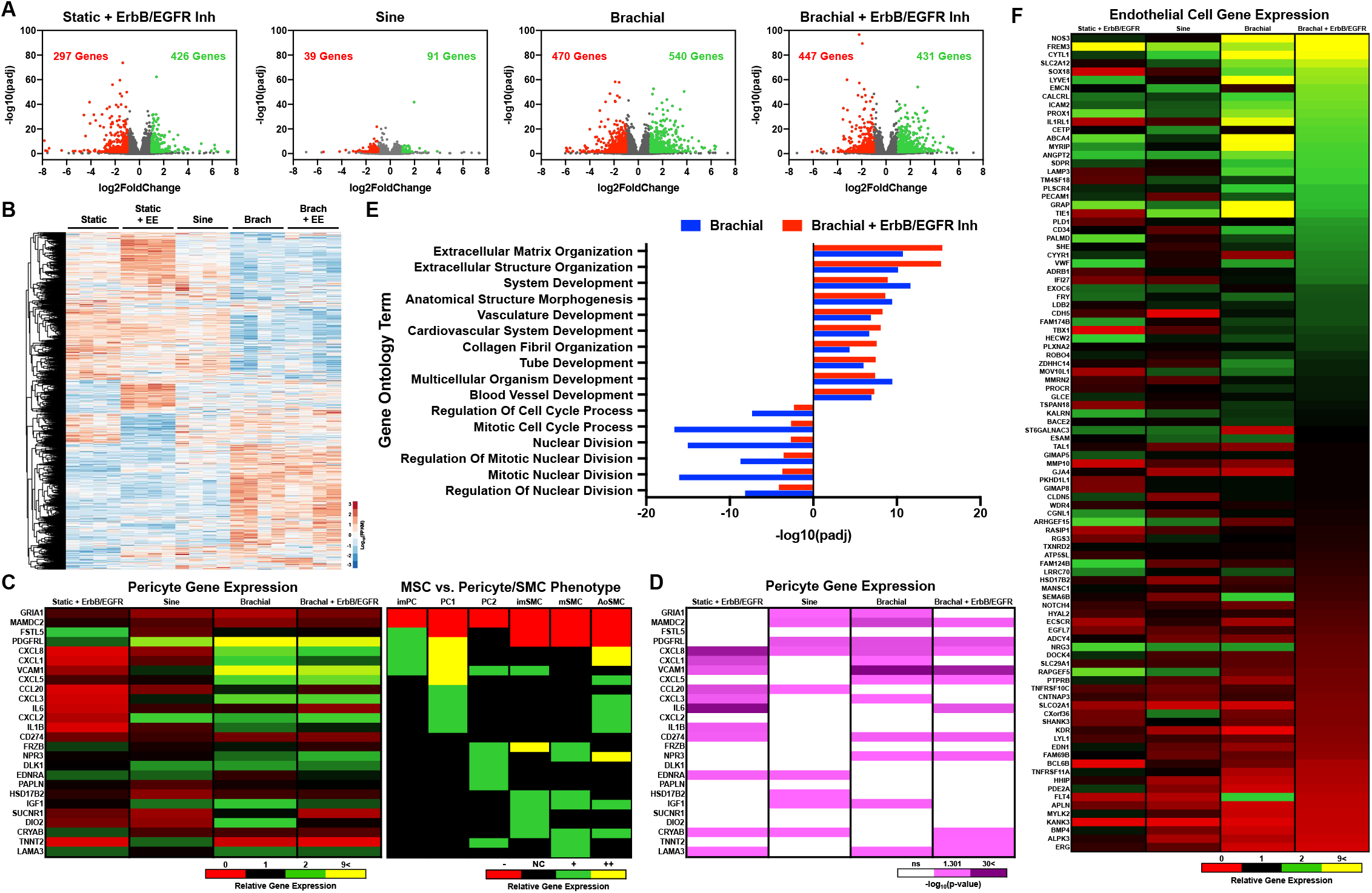
Gene expression analysis the RNAseq demonstrates that mechanical conditioning with brachial waveform loading enhances pericyte and endothelial cell gene expression. MSCs were treated with mechanical load and/or an ErbB/EGFR inhibitor for seven days. (A) Volcano plots of differentiation gene expression in comparison to the static control group. (B) Clustering analysis of the gene expression in the MSCs for significantly regulated genes. (C) The left heat map shows expression of genes for pericyte or vascular smooth cells in the treated cells. The right heat map is a visualization of how these genes change in bone marrow MSCs when they differentiate into mural phenotypes.^49^ The cell phenotypes listed are as follows: immature pericytes (ImPC), type 1 pericytes (PC1), type 2 pericytes (PC2), immature vascular smooth muscle cells (ImSMC), mature vascular smooth muscle cells (mSMC), and aortic vascular smooth muscle cells (AoSMC). (D) Statistical significance for the pericyte/mural cell related genes. (F) Expression of genes related to endothelial cell phenotype in the MSCs after mechanical and pharmacological conditioning.

### Optimal mechanical and pharmacological conditioning of MSCs increases pro-angiogenic potential *in vivo*

We next examined the effects of mechanical loading and pharmacological conditioning on the *in vivo* angiogenic properties of MSCs. We conditioned the cells with the various treatments for seven days and then implanted them subcutaneously in nu/nu mice in Matrigel. After 14 days, there were increased numbers of vessels invading the gels implanted with MSCs exposed to brachial waveform mechanical loading based on the macroscopic images of the implant (**Fig. 6A, B**). In particular, the cells treated with the brachial waveform loading and pharmacological co-treatment had the highest number of large vessels invading. Analysis with laser speckle imaging revealed increased perfusion in the skin over the implants of the groups treated with the brachial waveform loaded MSCs with pharmacological co-treatment, consistent with the macroscopic appearance of the implants (**Fig. 6C, D**). Histological analysis of the gels also demonstrated increased cells positive for PECAM, α-SMA and double positive for PECAM/α-SMA in those treated with brachial waveform loading in combination with pharmacological inhibitor treatment (**Fig. 6E, F**). In addition, there were increased nestin-positive cells with brachial loading and co-treatment with the E/E inhibitor as well as cells double positive for nestin and CD146 (**Fig. 6G, H**). There were also marked increases in PRGFRβ and PECAM/PDGFRβ positive cells with brachial loading and co-treatment with the E/E inhibitor (**Supplemental 18A, B**). We assessed proliferating cells in the Matrigel using staining for Ki-67 and found that only the static cells with the co-treatment E/E inhibitor had increased numbers of proliferating cells (**Supplemental Fig. 18C, D**). To confirm that the cells in the Matrigel were primarily derived from the MSCs that were delivered, we performed fluorescence ***in situ*** hybridization (FISH) for the human X chromosome (**Supplemental Fig. 19**). Together, these findings support the concept that combined conditioning with pharmacological inhibitors and specific mechanical conditions can increase the pro-angiogenic properties of MSCs.

**Figure 6.**
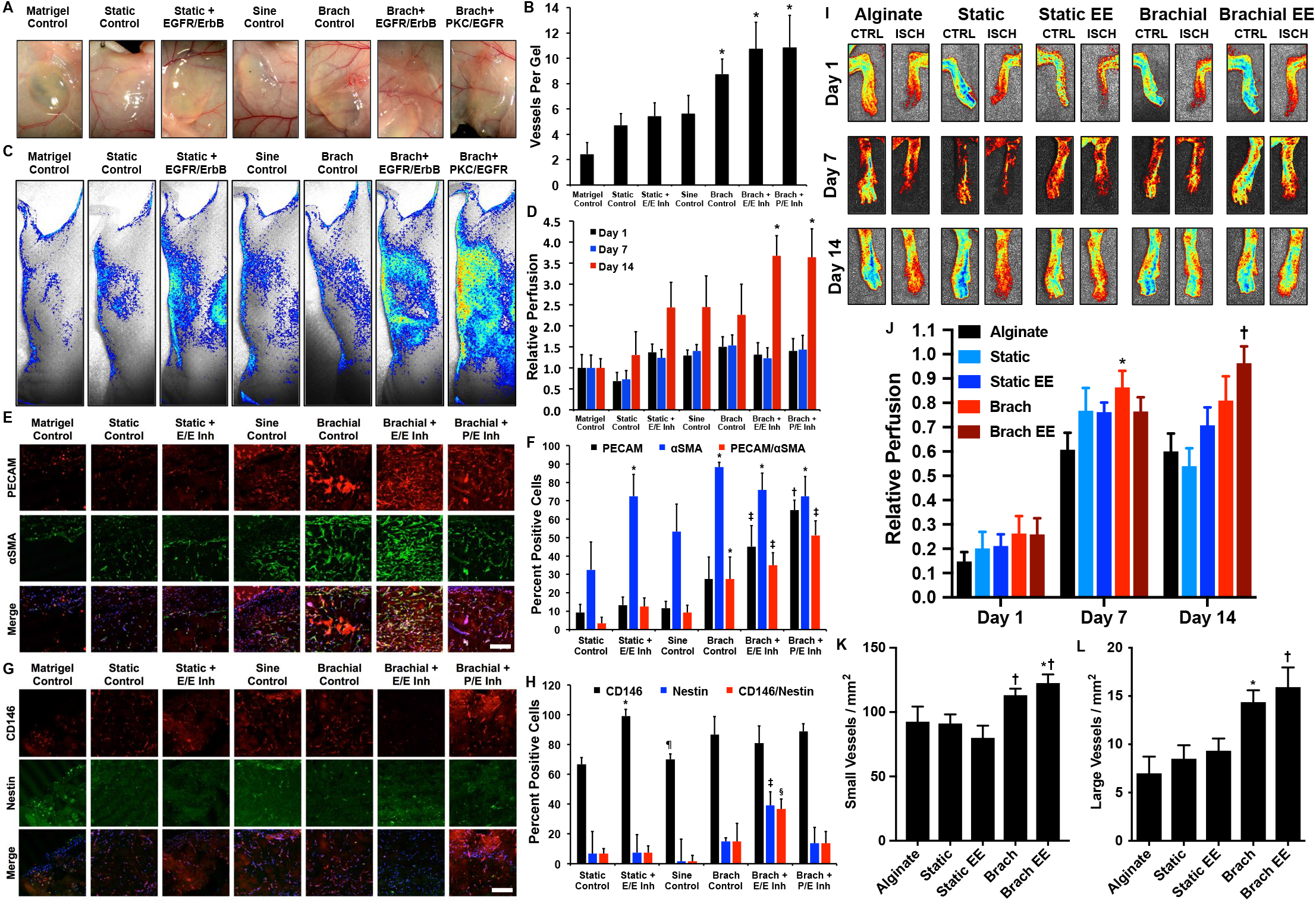
Optimized mechanical and pharmacological conditioning of MSCs increases their ability to induce angiogenesis and arteriogenesis following implantation subcutaneously or in a hind limb ischemia model. The MSCs were treated under the indicated conditions for seven days in culture and then implanted subcutaneously in nu/nu mice in Matrigel. (A) Photographs of the implants after 14 days of implantation. (B) Quantification of blood vessels in the gel using macroscopic images of the gel. *\*p* < 0.05 versus Matrigel, static, static with EGFR/ErbB inhibitor and sine groups. (C) Images of laser speckle imaging of the mice after 14 days of implantation. (D) Quantification of perfusion measured by laser speckle imaging following implantation. *\*p* < 0.05 versus Matrigel and static control groups. (E) Images of tissue sections from the gel regions of the explanted tissues immunostained for PECAM and α-SMA. Scale bar = 100 μm. (F) Quantification of the percent positive cells for the indicated markers. *\*p* < 0.05 versus static control groups. †*p* < 0.05 versus static, static with EGFR/ErbB inhibitor, sine and brachial groups. ‡*p* < 0.05 versus static, static with EGFR/ErbB inhibitor and sine groups. (G) Images of tissue sections immunostained for CD146 and Nestin. (H) Quantification of the percent positive cells for the indicated markers. *\*p* < 0.05 versus static control groups. †*p* < 0.05 versus static, static with EGFR/ErbB inhibitor, sine and brachial groups. ‡*p* < 0.05 versus static, static with EGFR/ErbB inhibitor and sine groups. §*p* < 0.05 versus static and sine groups. ¶*p* < 0.05 versus static with EGFR/ErbB inhibitor group. Scale bar =100 μm. (I) Laser speckle imaging of the feet of mice implanted with MSCs treated for seven days with the indicated treatments. (J) Quantification of the perfusion in the feet of the mice after induction of hind limb ischemia. \**p* < 0.05 versus alginate group. †*p* < 0.05 versus alginate and static groups. (K) Quantification of the number of small blood vessels in the thigh muscle of the ischemic limb in the mice implanted with MSCs conditioned with the treatments. \**p* < 0.05 versus the static groups. †*p* < 0.05 versus the static EE group. (L) Quantification of large vessels in the thigh muscle of the ischemic limb of the mice. \**p* < 0.05 versus the alginate group. †*p* < 0.05 versus the alginate, static and brachial EE groups.

To further confirm the functional activity of the conditioned MSCs, we exposed the cells to the treatments for seven days, encapsulated them in alginate-RGD/collagen gel and implanted them in nu/nu mice that underwent surgery to create unilaterial hindlimb ischemia through femoral artery ligation. We observed enhanced recovery of perfusion in the mice treated with cells that had been conditioned with brachial waveform loading and E/E inhibitor (**Fig. 6I, J**). Histological analysis revealed a moderate increase in capillaries and an over two-fold increases in larger mature vessels in the muscle from mice treated with cells conditioned with brachial loading or brachial loading with E/E inhibitor treatment (**Fig. 6K, L; Supplemental Fig. 20**). We confirmed the continued presence of MSCs in the hind limb of the mice using FISH for the human X chromosome (**Supplemental Fig. 21**).

## Discussion

Mesenchymal stem cells are an appealing cell type for use in therapies for ischemia but clinical trials have not yielded consistent benefits for patients. The application of MSCs in vascular regeneration and tissue engineering has been limited by contradictory or inconsistent findings in studies describing MSC differentiation into vascular phenotypes^21,24^. Our studies identified conditions that enhanced MSC therapeutic activity and induced increased differentiation toward a novel phenotype that had combined markers of pericytes and endothelial cells. There have been several studies that support that perivascular cells may behave as MSCs in the body^42^. However, other studies suggest that pericytes are not stem cells^43^. Our study demonstrates that the pericyte phenotype of MSCs can be increased by conditioning with mechanical load and, further, that enhancing this hybrid phenotype can improve the therapeutic potential of these cells.

In our mechanobiological screen, we maximized the signaling through the p-Smad2/3 and Yap/Taz pathways. These pathways have been linked to MSC differentiation into other MSC phenotypes^53^. However, these pathways have also been linked to angiogenesis and pericyte function. Signaling through Yap/Taz is enhanced in the tip cells of capillaries during angiogenesis and vascular development^54–56^. In addition, pericyte-specific knockout of Yap and Taz disrupts their coordination of alveologenesis by reducing their production of HGF^57^. Signaling through Smad2/3 is required for vascular stability^58^, modulates pericyte/endothelial interactions^59^ and TGF-β signaling enhances the association of pericytes/mesenchymal cells to endothelium cells^60^. There is also evidence that links Smad2/3 to endothelial differentiation in MSCs and induced pluripotent stem cells^61,62^. In our study, optimal conditions to maximize these pathways enhanced the pericyte and endothelial cell-like phenotype of the MSCs. In addition, these treatments enhanced the expression of HGF and the angiogenic potential of the soluble factors produced by the MSCs. The enhanced pericyte/endothelial cell phenotype of the mechanical/biochemically conditioned MSCs was supported by expression of FACS markers, increased tube formation alone or in co-culture with endothelial cells, increased production angiogenic soluble factors, gene expression by RNAseq and enhanced angiogenic activity after subcutaneous implantation or in a hind limb ischemia model.

Previous studies have identified that mechanical forces can increase the vascular cell phenotype of MSCs. Notably, the application of mechanical strain to MSCs increases the expression of vSMC markers^63,64^ and the application of shear stress has in some studies been linked to the expression of endothelial cell markers^65^. There have also been mixed results in studies aiming to induce endothelial differentiation in MSCs, with some studies finding differentiation into endothelial phenotype with VEGF-A treatment while others do not show these effects^21,24,66^. Our findings using flow cytometry suggest that there very little pure endothelial cell differentiation of MSCs under a broad range of treatments. Given our findings of a pericyte/endothelial hybrid phenotype, even the presence of multiple endothelial markers does not exclude the possibility the MSCs also express mural markers as well. Even under the most optimal conditions, only around eight percent of the total MSC population expressed endothelial markers in the absence of pericyte markers. Thus, our studies suggest that previous findings of endothelial differentiation in MSCs should be viewed with the perspective that further studies may be needed to rule out hybrid phenotypes.

In summary, our findings have identified that MSCs can be conditioned into an enhanced phenotype with increased expression of markers of pericytes and endothelial cells. The conditioning required the use of complex mechanical strain waveforms and drug treatment. We were not able to achieve maximal regenerative properties from the MSCs in the absence of applied mechanical forces. Given the wide variation in MSC behavior and loss of activity due to disease and aging, a potential approach may be to first isolate MSCs from the patient and optimize the desired phenotype using a mechanobiological screen similar to the one used in this study. For vascular regeneration, we envision that MSCs could be harvested from patients and tested in a mechanobiological screen to maximize the mixed EC/pericyte population. These mechanical and biochemical/pharmacologically optimized and conditioned cells can then be implanted or injected into the patient for treatment. Thus, while the pathways and mechanical conditions that were found to be optimal in this study may be generalizable to many MSC lines, the overall optimization process of using mechanobiological screening would likely provide a more robust method for enhancing stem cell therapies.

## Acknowledgements

The authors gratefully acknowledge funding through the American Heart Association (17IRG33410888), the DOD CDMRP (W81XWH-16-1-0580; W81XWH-16-1-0582) and the National Institutes of Health (1R21EB023551-01; 1R21EB024147-01A1; 1R01HL141761-01) to ABB.

## Materials and Methods (Online)

### Construction of a high throughput biaxial oscillatory strain system

Mechanical strain was applied to cell culture using a high throughput system described previously^34,36,67^. Briefly, cells are cultured on custom made plates that are mounted on a system that applies strain. The cell culture plates comprised of a 0.005” thin silicone membranes (Specialty Manufacturing, Inc.) that are sandwiched between two plates, with silicone rubber gaskets at the interfaces to prevent leaking. These cell culture membranes are UV sterilized and coated with 50 μg/mL fibronectin overnight at 37°C to allow cell adhesion. After cell attachment, the plates are mounted on to the top plate of the system using screws. To apply mechanical strain, a platen with 576 Teflon pistons is moved into the cell culture membrane. The motion is driven by a hygienically sealed, voice coil-type linear motor (Copley Controls). The platen is stabilized using six motion rails mounted with linear motion bearings. The hygienically sealed motor housing has chilled water running through in order to prevent over heating during operation. The 576 pistons that come in contact with the cell culture surface can be individually adjusted to have different height, allowing precise calibration of the strain applied to each well.

### Cell lines and cell culture

Human mesenchymal stem cells (Millipore, Inc.) were cultured in low glucose DMEM medium supplemented with 15% fetal bovine serum, L-glutamine and penicillin/streptomycin. Following trypsinization, cells that reached passage 6 were seeded on the membranes at 20,000 cells per cm^2^ before mechanical loading. Human umbilical vein endothelial cell (HUVECs; PromoCell GmbH) were cultured in MCDB 131 medium supplemented with 7.5% fetal bovine serum, L-glutamine and penicillin/streptomycin and EGM-2 SingleQuot Kit (Thermo Fisher Scientific, Inc.). All cells were cultured in an incubator at 37 °C under a 5% CO_2_ atmosphere.

### Measurement of transcription factor activity

Human mesenchymal stem cells (passage 3) were transduced with lentiviruses for the expression of luciferase reporter constructs (Qiagen) for the target transcription factors. Briefly, cells were cultured with the lentivirus (1 x 10^7^ TU) in media containing polybrene for 24 hours. Following virus incubation, media was replaced with normal media for a day. Transduced cells were then selected with puromycin (1 μg/mL) containing media for three days. Following the treatments, the cells were lysed and the relative luminescent signal was measured using the luciferase assay kit (Promega) and read with the FlexStation-3 plate reader (Molecular Devices). For each plate, average from three individual readings was reported.

### Immunocytochemical staining

Following the treatments, the cells were fixed in 4% paraformaldehyde in PBS for 10 minutes followed by washing and permeabilization with 0.1% Triton X-100 PBS for 5 minutes. Next, samples were blocked with PBS containing 5% FBS and 1% BSA for 40 minutes. After washing, cells were incubated with primary antibodies (see **Supplemental Table 1** for specific antibodies and concentrations) in PBS with 1% BSA overnight at 4°C. The samples were then washed twice in PBS with 1% BSA and incubated with secondary antibodies in PBS with 1% BSA for 2 hours in a light protected environment. For actin staining, the samples were treated with Alexa 594 conjugated phalloidin for 30 minutes. Cells treated with extensive washes with PBS with 1% BSA prior to mounting in anti-fade media (Vector Laboratories, Inc.). The samples were then imaged using epifluorescence microscopy (Axio Observer; Carl Zeiss, Inc.), or confocal microscopy using either an LSM 710 laser scanning confocal microscope (Carl Zeiss, Inc.) or an FV10i Confocal Laser Scanning Microscope (Olympus, Inc.).

### Cell lysis and immunoblotting

Following the treatments, the cells were lysed in 20 mM Tris with 150 mM NaCl, 1% Triton X-100, 0.1% SDS, 2 mM sodium orthovandate, 2 mM PMSF, 50 mM NaF and a protease inhibitor cocktail (Roche, Inc.). The proteins were separated on a NuPAGE 10% bis-tris midi gel (Novex) and transferred to nitrocellulose membrane using iBlot transfer stack (Novex). The membranes were blocked for one hour in 5% non-fat milk in PBS with 0.01% tween-20 (PBST). After washing twice in PBST, cells were incubated with primary antibodies (**Supplemental Table 2**) overnight in 1% non-fat milk at 4°C. The membranes were washed with PBST and incubated at room temperature for two hours with secondary antibody. The membrane was treated with chemiluminescent substrate (SuperSignal West Femto; Thermo Fisher Scientific, Inc.) then imaged using a digital imaging system (Cell Biosciences, Inc.).

### Long term conditioning of hMSCs using biochemical factors

For long-term conditioning, cells were incubated with the treatments as shown in **Supplemental Table 3**. The cells were treated with mechanical loading for 4 hours/day for 7 days under sine and brachial waveform at 0.1 Hz and 7.5% maximum strain, or cultured under static conditions. The culture media containing the treatments were replaced on day 3 and day 5 for all treatments except 5-Aza. Cells that were treated with 5-Aza had their culture media replaced after 24 hours with standard media for the rest of the experiment. In some cases, culture media was replaced with 0.5% FBS on the final day of loading to allow the harvest of conditioned media without the presence of the treatments.

### Flow Cytometry

For the separation of cells by markers of vascular phenotypes, the cells were detached from plate using Accutase (Sigma-Aldrich) and were labeled with fluorescent antibodies according the BD Bioscience flow cytometry staining kit protocol (BD 562725; see **Supplemental Table 4** for the specific antibodies used). Briefly, the detached cells were centrifuged and the supernatant was removed. Fixing and permeabilizing buffer was added while the cells were vortexed and incubated for 40 min. Next, the samples were centrifuged and the supernatant was removed. Cells were then treated with washing buffer containing antibodies for 50 min. Following antibody incubation, cells were centrifuged with washing buffer two more times, before they are treated with stain buffer and measured. A BD LSR II Fortessa Flow Cytometer (BD Biosciences) was used to measure population fluorescent signals. At least 10,000 events were recorded and further gating and quantification was done through FlowJo software.

### In vitro tube formation assay

A day prior to tubule formation assay, growth factor reduced Matrigel (Corning, Inc.) was allowed to thaw overnight at 4°C. Human umbilical cord endothelial cells (HUVECs; passage 5) were labeled with CellTracker Red CMTPX Dye (Thermo Fisher Scientific, Inc.) and hMSCs were labeled with cell tracker green (Thermo Fisher Scientific, Inc.). These cells were then cultured in 0.5% FBS containing media for 16 hours prior to the tubule formation assay. On the day of the assay, glass bottom 96-well plates were coated with 50 μl of Matrigel per well and then incubated for 30 minutes at 37°C. The fluorescently labeled cells were passaged with the conditioned media from the long term loading onto the plates at a total seeding density of 20,000 cells/well in either hMSC alone, HUVEC alone, or a co-culture of both hMSC and HUVEC at 1:1 ratio. These cells were then imaged at 0 hour, 10 hours and 22 hours post seeding using Cytation 5 Cell Imaging Multi-mode Reader (Biotek) in the facilities of the Targeted Therapeutic Drug Discovery & Development Program (TTP) at UT Austin. For quantification, the number of complete loops formed was counted.

### Measurement of soluble factor production

Conditioned media was assayed for soluble factor production using an antibody array for angiogenic factors (Proteome Profiler Human Angiogenesis Array Kit; R&D Systems, Inc.). In addition, the concentrations of some of the factors were measured using ELISA assays per manufacturer’s instructions (R&D Systems, Inc.).

### Subcutaneous implantation in nu/nu mice

All animal studies were performed with the approval of the University of Texas at Austin Institutional Animal Care and Use Committee (IACUC) and in accordance with NIH guidelines “Guide for Care and Use of Laboratory Animals” for animal care. To assess the *in vivo* response of conditioned hMSCs, the cells were implanted subcutaneously in nu/nu mice. Following seven days of conditioning with the treatments, the cells were detached using 0.05% Trypsin-EDTA and spun down at 200 g for three minutes. The supernatant was removed and the cells were resuspended in Matrigel (Corning) at 2×10^6^ cells/mL, in a total volume of 200 μL. These cell suspensions were then injected subcutaneously on the dorsal surface of the six-week old nu/nu mice (Jackson Laboratories, Inc.). Blood perfusion on the back of the mice was assessed using a custom speckle imaging system on 1, 3, 5, 7 and 14 days following implantation of cells^68^. Laser speckle imaging was quantified by taking the ratio of the perfusion in the region of skin over the implanted Matrigel to the perfusion of the skin over the sacral region of the mouse. After 14 days, the mice were euthanized and the tissue harvested for histological analysis. All animal studies were performed with the approval of the University of Texas at Austin Institutional Animal Care and Use Committee and in accordance with NIH guidelines “Guide for Care and Use of Laboratory Animals” for animal care.

### Histochemical staining and immunohistochemistry of the skin tissues

Tissues from the subcutaneous study were cryopreserved. The subcutaneous Matrigel plug was excised using a 10 mm sterile biopsy punch. The tissues were fixed in 10% formalin in PBS for 24 hours. Next, the fixed tissues were submerged in 30% sucrose in PBS for 4 days. Tissue samples were then frozen in isopentane cooled with liquid nitrogen. Frozen tissue samples were sectioned to create 20 μm thick sections. Briefly, the frozen tissue slices were incubatd in PBS for 5 min. The samples were then incubated with Fc receptor blocker (Innovex Biosciences) for 30 minutes and then blocked with 25% FBS in PBS for 45 min. After two washes with 1% BSA in PBS, samples were incubated with antibodies found in **Supplemental Table 5** overnight followed by secondary antibody staining for 2 hours. Samples were then mounted with Vectashield and imaged using FV10i Confocal Laser Scanning Microscope (Olympus, Inc.). For H&E staining, the frozen tissues were sectioned at 10 μm prior to staining.

### Cell encapsulation in alginate beads

The hMSCs at passage 5 were mechanically conditioned with either static, sinusoidal or brachial strain waveform at 0.1 Hz, 7.5% maximal strain for 7 days. During loading, the cells were treated with either no treatment or 1 μM EGFR/ErbB inhibitor biomolecule. The conditioned cells were then detached using 0.05% EDTA trypsin, spun down, and the supernatant removed. Approximately 1×10^6^ cells were then resuspended in 200 μL alginate solution with 2% RGD peptides and 0.045% collagen. The mixture was extruded to form alginate beads with bead diameter of 1200 μm, which were then applied to an ischemic leg of the mice.

### Hind limb ischemia model in nu/nu mice

To assess the angiogenic potential and the functional recovery induced by the MSCs, we performed a hind limb ischemia mice model in nu/nu mice (Jackson Laboratories, Inc.). At ten-weeks of age, the left femoral artery and branches were ligated to induce ischemia. Conditioned cells were encapsulated in alginate beads as described above and then implanted onto the ischemic leg. Blood flow through the limb was assessed using a speckle laser imaging system on day 1, 3, 5, 7, and 14 days. The ratio between the flow though the ischemic limb to the control limb was measured to assess recovery from ischemia with correction for illumination intensity. After 14 days, the tissues were fixed using formalin perfusion and muscles of the hind limbs harvested.

### RNA Sequencing and Analysis

Following treatments, RNA was isolated from the cells using the Qiagen RNeasy^®^ Mini Kit. RNAseq was performed using a Illumina HiSeq 4000. For sequencing, single reads of 50 base pairs were performed after poly-A mRNA capture using the Poly(A) Tailing Kit (Ambion) and Ultra II Directional RNA Library Prep Kit (NEB) to isolate mRNA and perform dUTP directional preparation of the mRNA library. RNA sequencing was performed by the Genomic Sequencing and Analysis Facility at UT Austin. Gene expression analysis was performed using R. Gene ontology was performed using the Molecular Signatures Database (En).

### Histochemical staining and immunohistochemistry of the skin tissues

Muscles from the ischemic tissues were first deparaffinized. Slides were then treated with DAKO antigen retrieval solution (DAKO) at 75 °C for 3 hours. Samples were then blocked with 25% FBS in PBS for 45 min. Slides were then stained for PECAM DAB marker from DAKO kit (Agilent Cat# K406511-2). Briefly, samples were blocked with dual enzyme blocker solution from DAKO. Slides were then incubated with PECAM-1 antibody (Abcam Cat#ab28364) overnight. On the following day, samples were labeled with HRP labed secondary antibody for 30 minutes followed by extensive washes with PBS. Slides were developed with DAB chromagen for 1 minute and then counterstained using Mayer’s hematoxylin.

### Fluorescence In Situ Hybridization (FISH)

Tissue slides were labeled with FISH probes following the kit instruction from VividFISH FFPE pretreatment and VividFISH CEP probe (Genecopoeia). Briefly, tissue slides were treated with pretreatment solution at 85 °C for 90 minutes followed up with a protease solution for 20 minutes. Slides were then denatured at 73 °C for 5 minutes, and treated with FISH probe mixed with the hybridization solution overnight at 42 °C. On the following day, samples were washed and treated with DAPI mounting medium. The slides were imaged using an FV10i Confocal Laser Scanning Microscope (Olympus, Inc.).

### Statistical analysis

All results are shown as mean ± standard error of the mean. Multiple comparisons between groups were analyzed by two-way ANOVA followed by a Tukey post-hoc or a Dunnett post-hoc test when testing multiple comparisons versus a control group. For non-parametric data, multiple comparisons were made using the Kruskal-Wallis test followed by post-hoc testing with the Conover-Iman procedure. A p-value of 0.05 or less was considered statistically significant for all tests.

## Supplementary Figure Legends

**Supplemental Figure 1.**
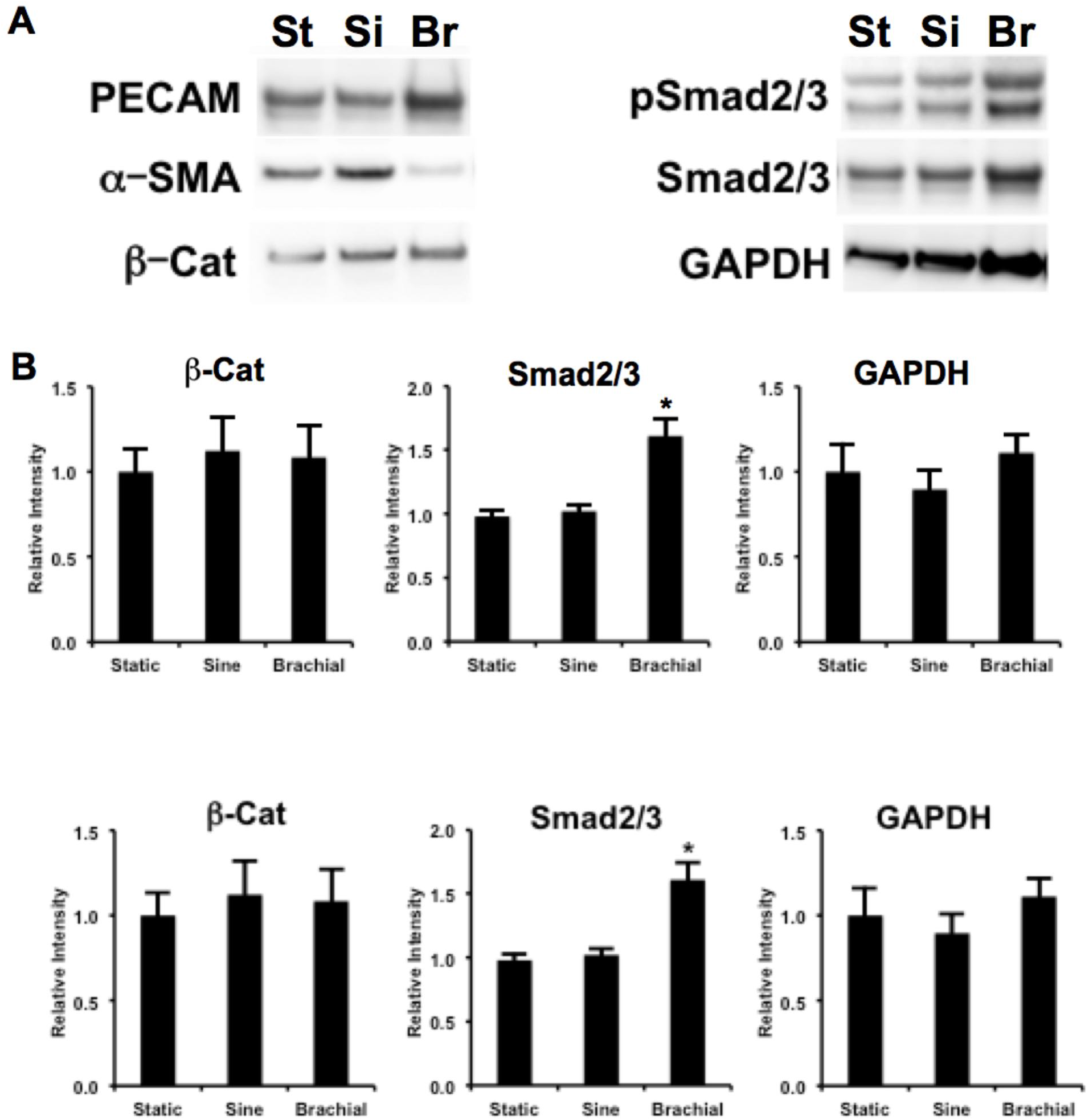
(A) Western blots on lysates from MSCs treated with mechanical loading for 8 hours at 7.5% strain. (B) Quantification of western blots from MSCs treated with mechanical loading.

**Supplemental Figure 2.**
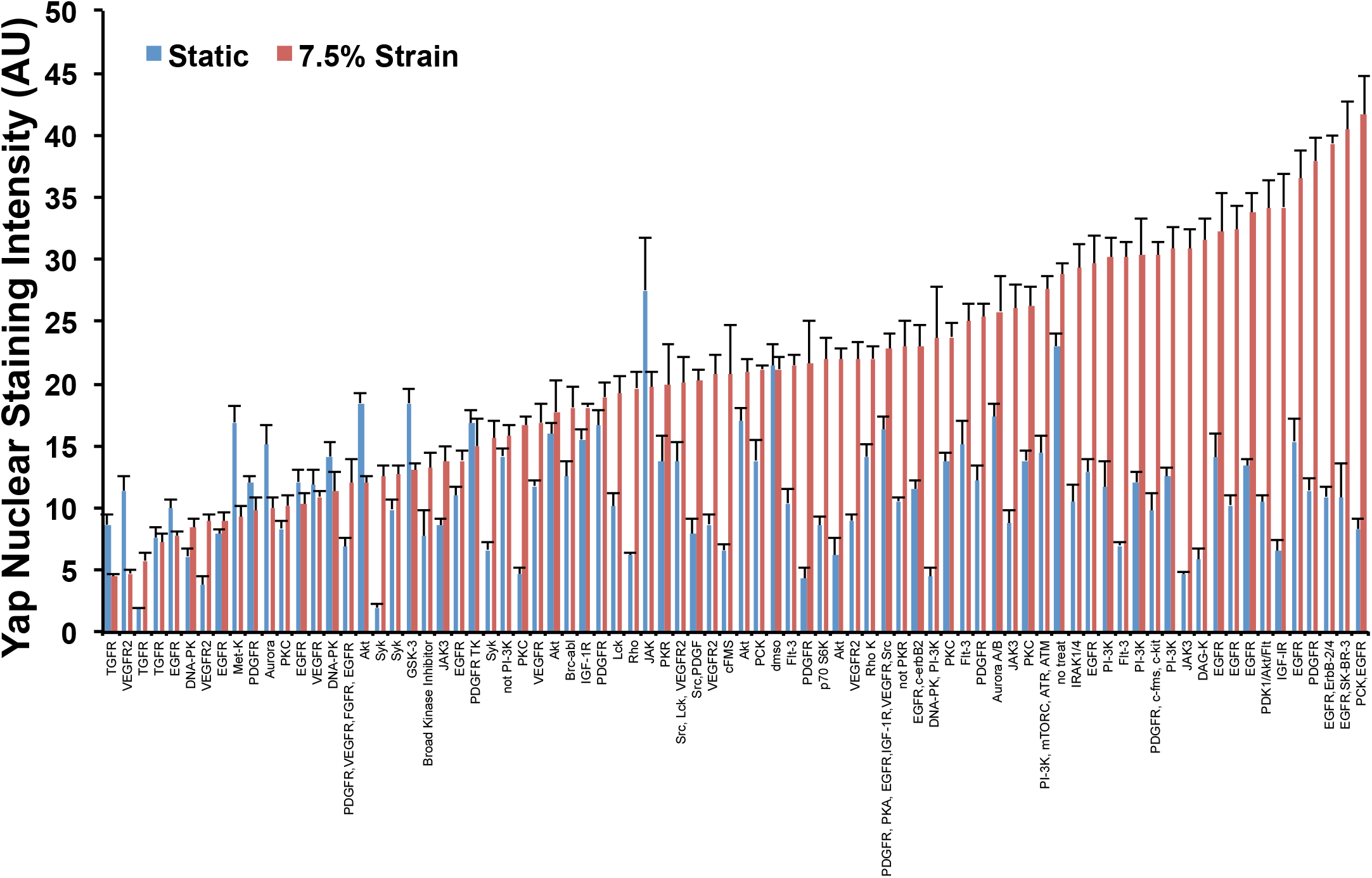
High throughput drug screen for Yap nuclear localization following seven days of loading with co-treatment with kinase inhibitors.

**Supplemental Figure 3.**
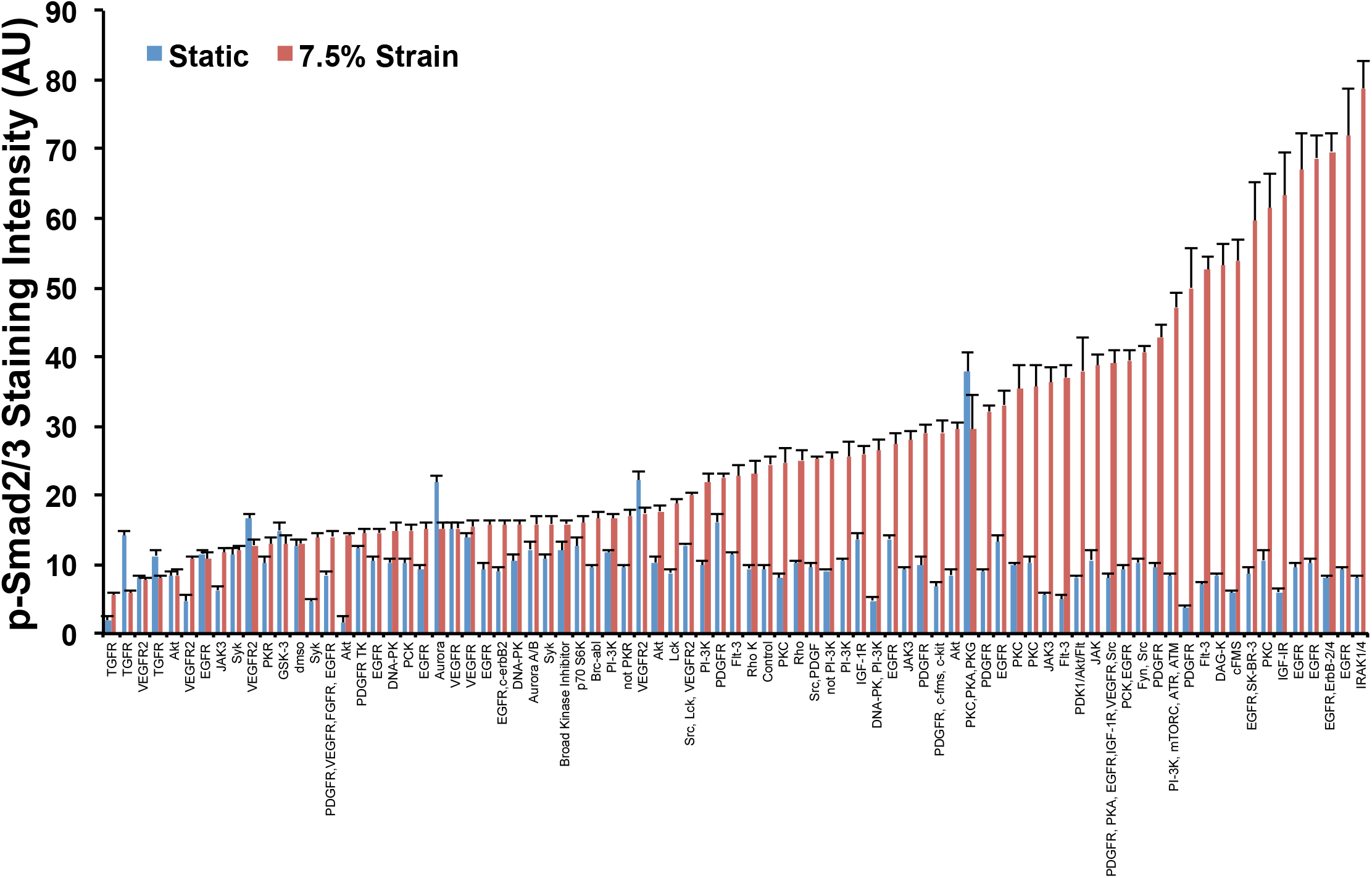
High throughput drug screen for p-Smad2/3 activation localization following seven days of loading with co-treatment with kinase inhibitors.

**Supplemental Figure 4.**
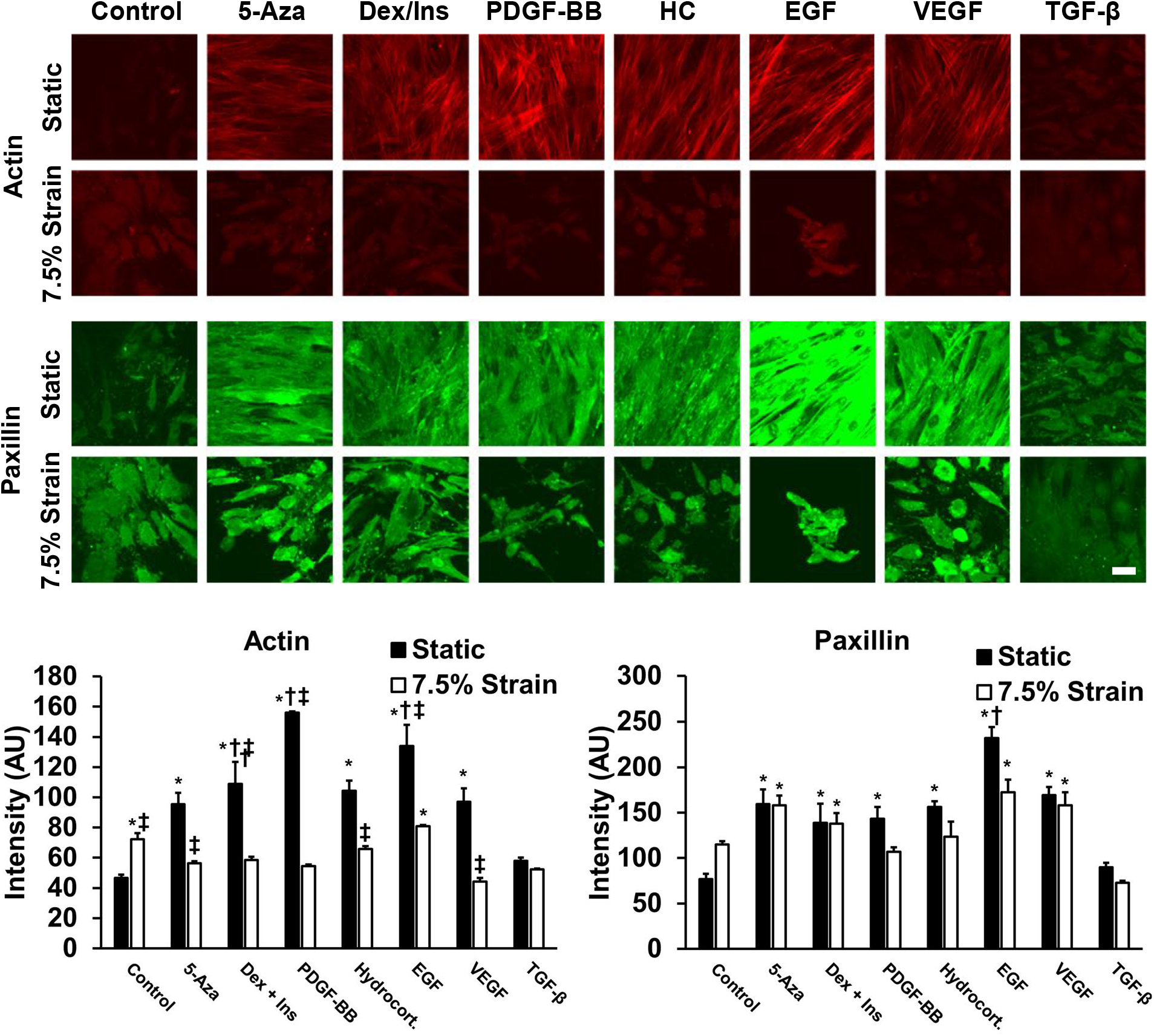
Staining for actin and paxillin after 7 days of loading at 7.5% maximum strain at 0.1 Hz. *\*p* < 0.05 versus cells treated with static conditions under baseline treatment. †*p* < 0.05 versus mechanically strained group under baseline treatment. ‡*p* < 0.05 versus cells under static conditions with the same biochemical treatment. Scale bar = 100 μm.

**Supplemental Figure 5.**
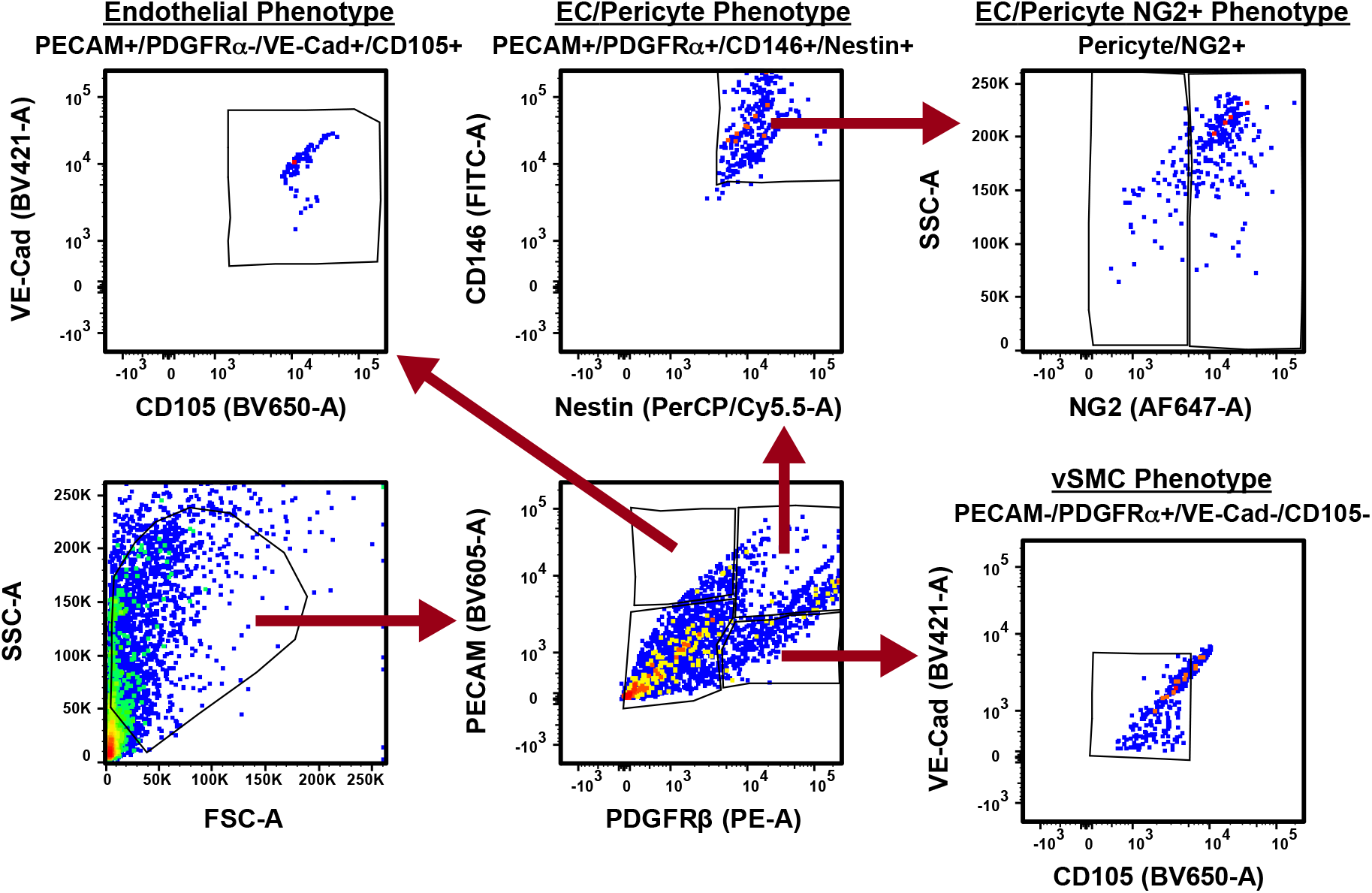
Gating strategy for flow cytometry analysis for vascular lineages in treated MSCs.

**Supplemental Figure 6.**
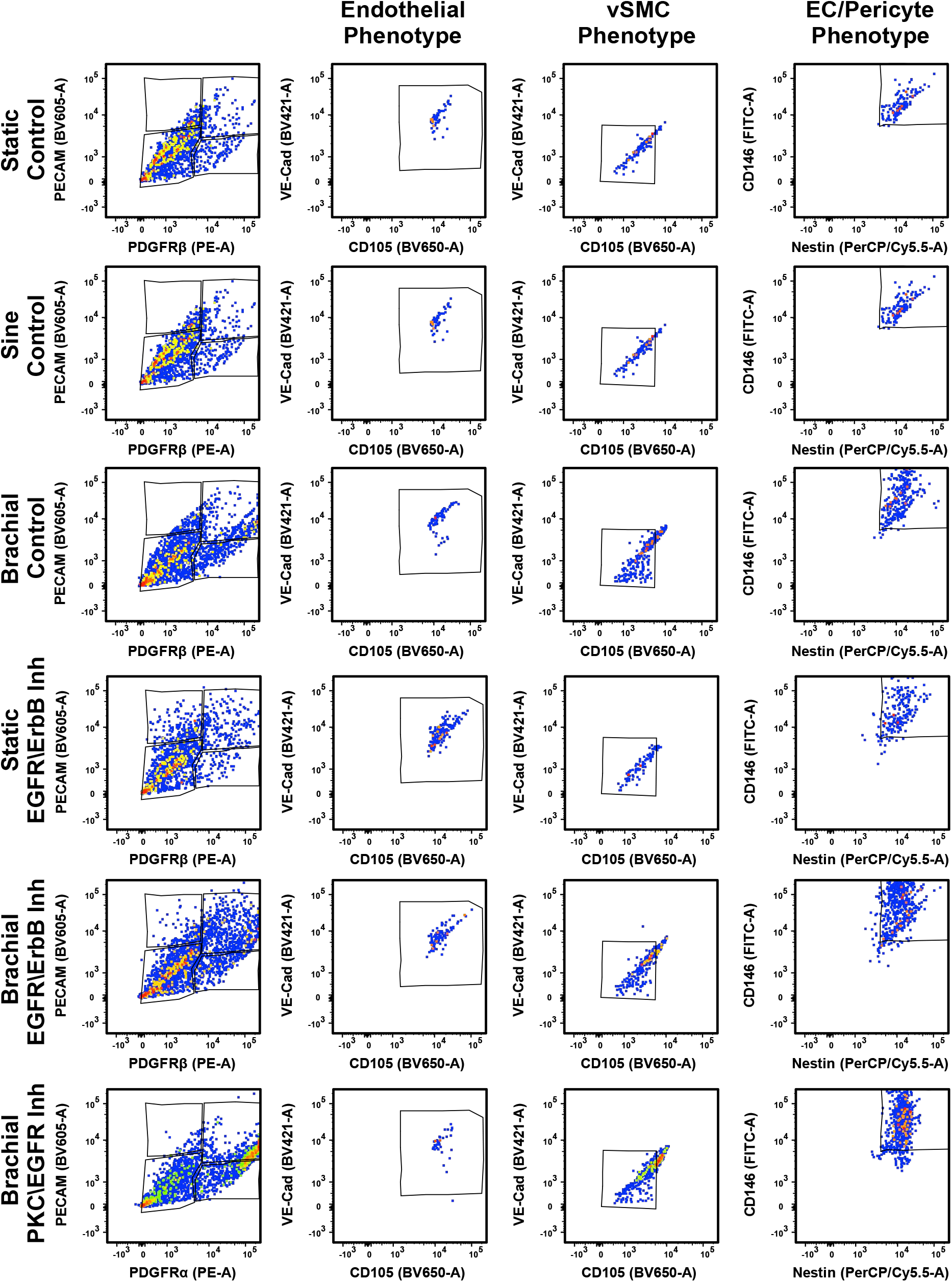
Representative separation of cell populations for the flow cytometry analysis.

**Supplemental Figure 7.**
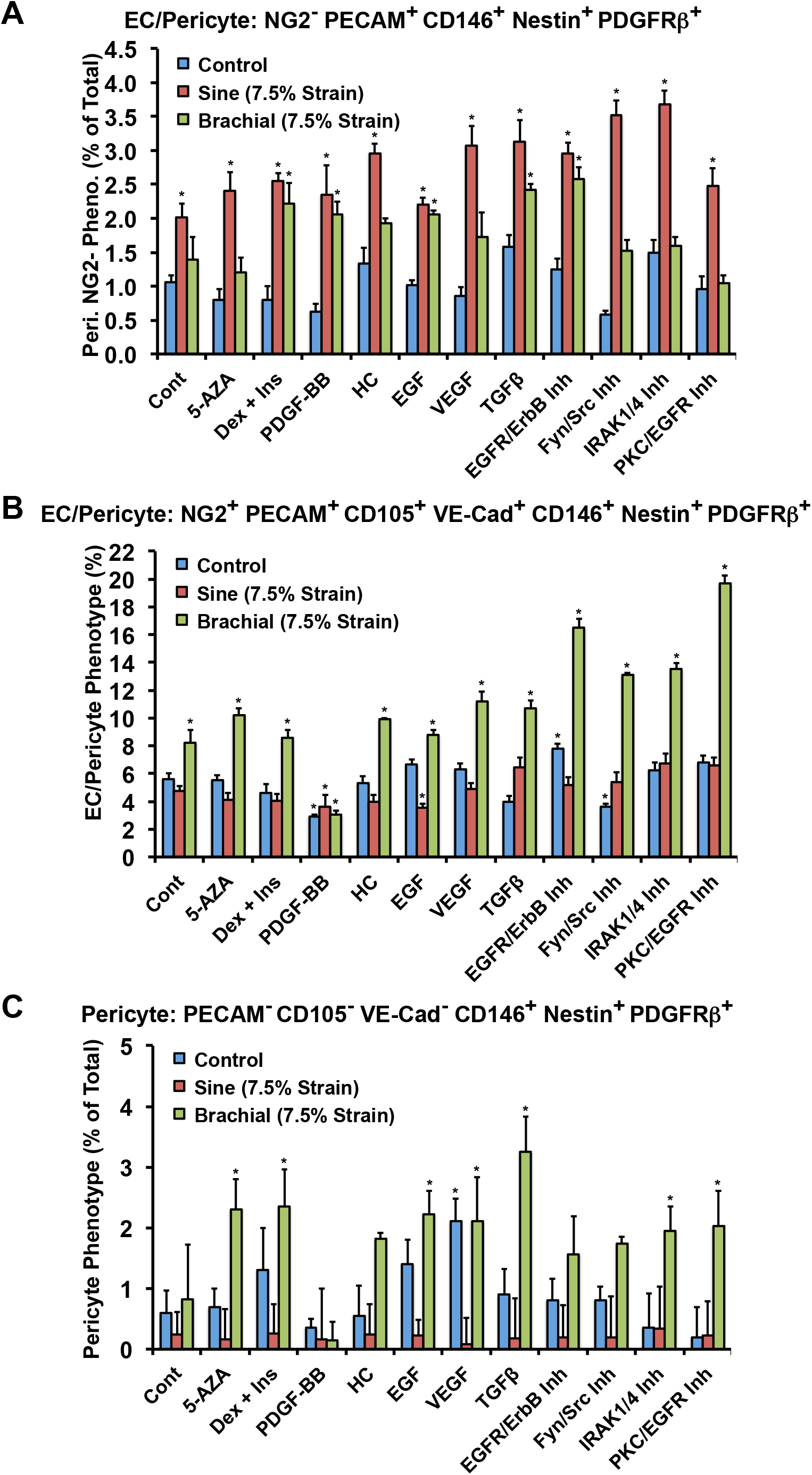
Quantification of cell populations for (A) endothelial cell/pericyte phenotype that are NG2^-^, (B) endothelial cell/pericyte phenotype that are also positive for additional endothelial markers and (C) pericyte phenotype. *\*p* < 0.05 versus MSCs under static control conditions.

**Supplemental Figure 8.**
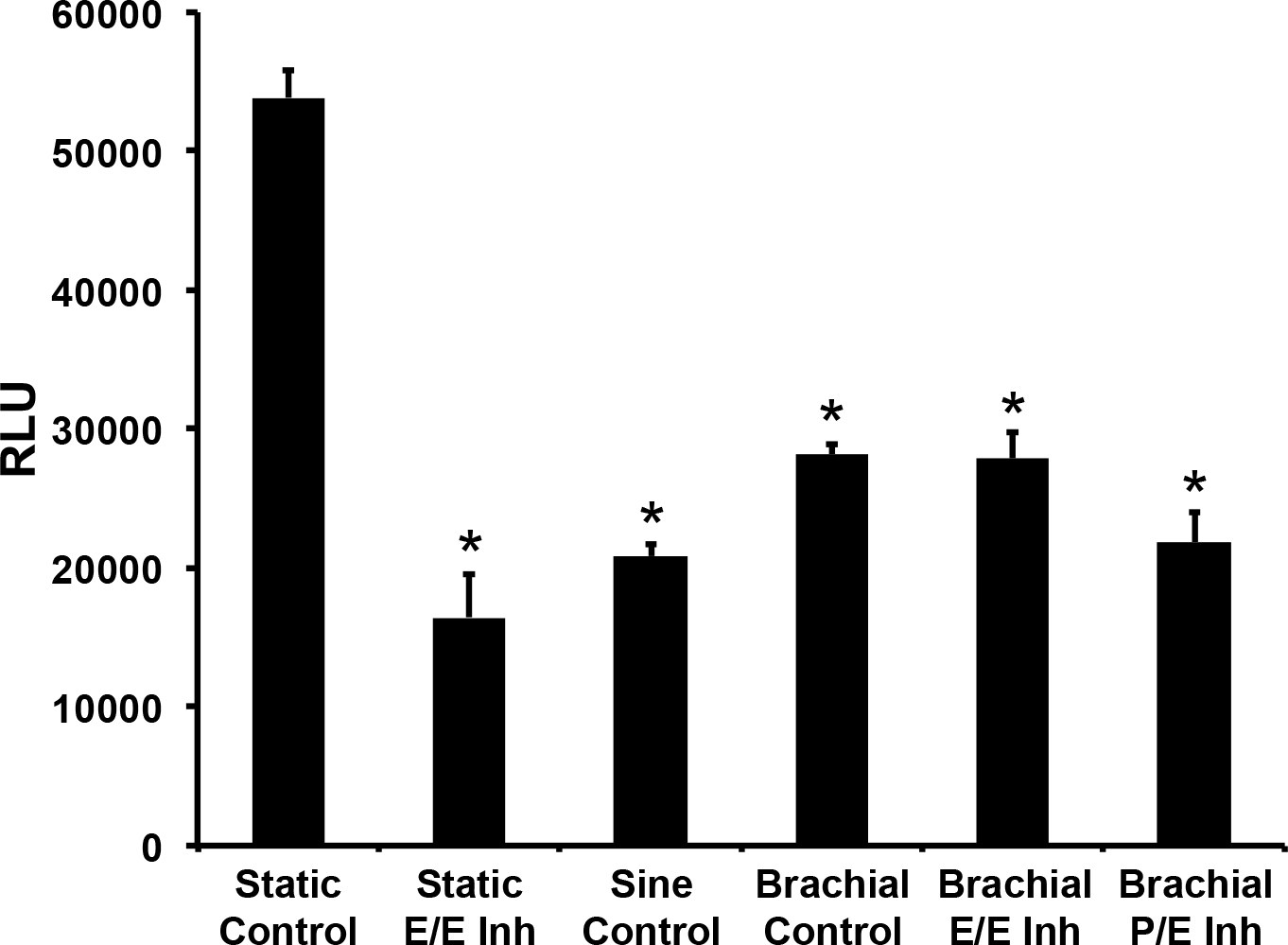
Proliferation of endothelial cells under treatment with conditioned media from MSCs treated with mechanical load and pharmacological inhibitors for seven days. \**p* < 0.05 versus static control group.

**Supplemental Figure 9.**
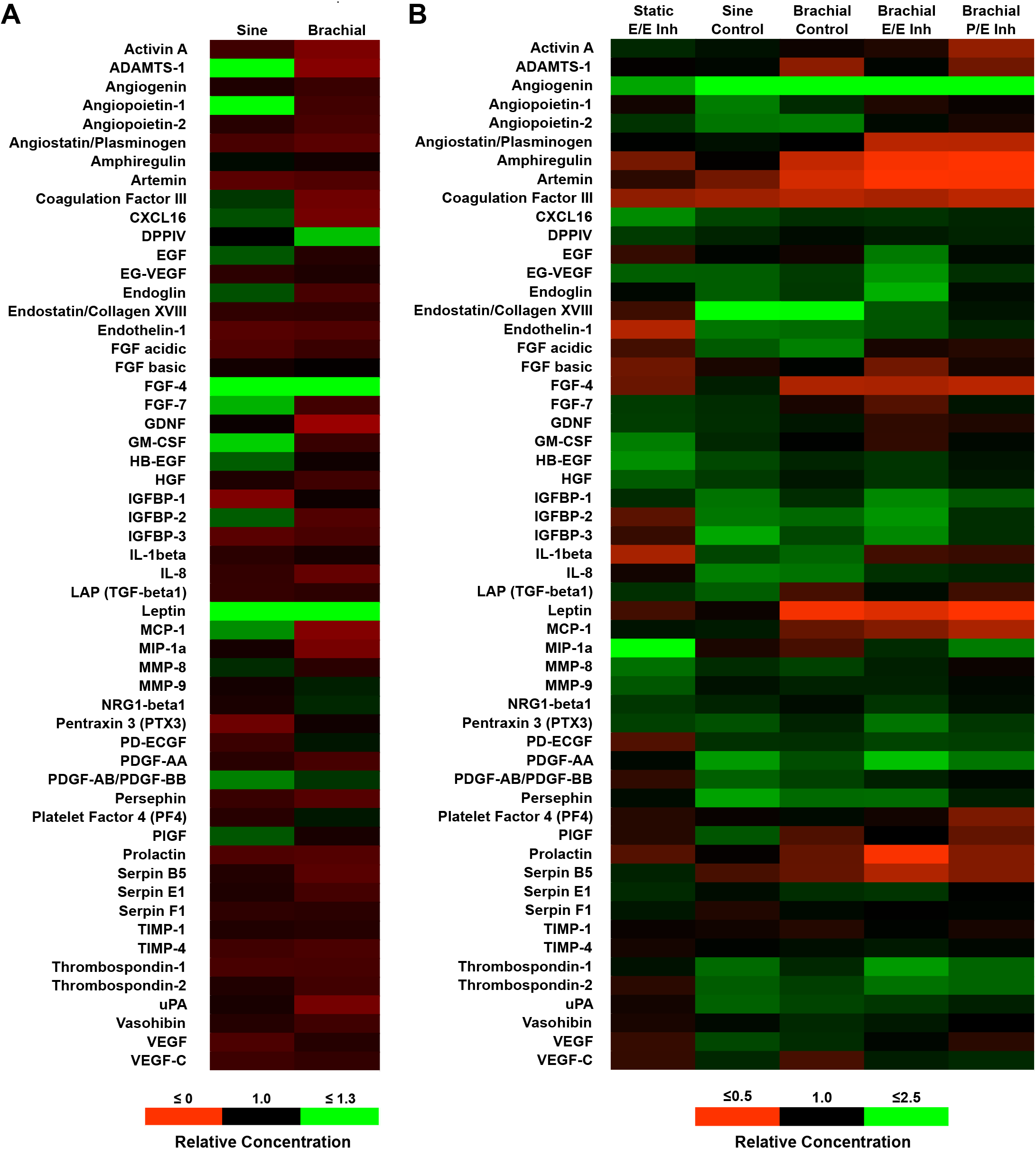
Antibody array results for conditioned media from MSCs treated with mechanical load for (A) 24 hours or (B) seven days.

**Supplemental Figure 10.**
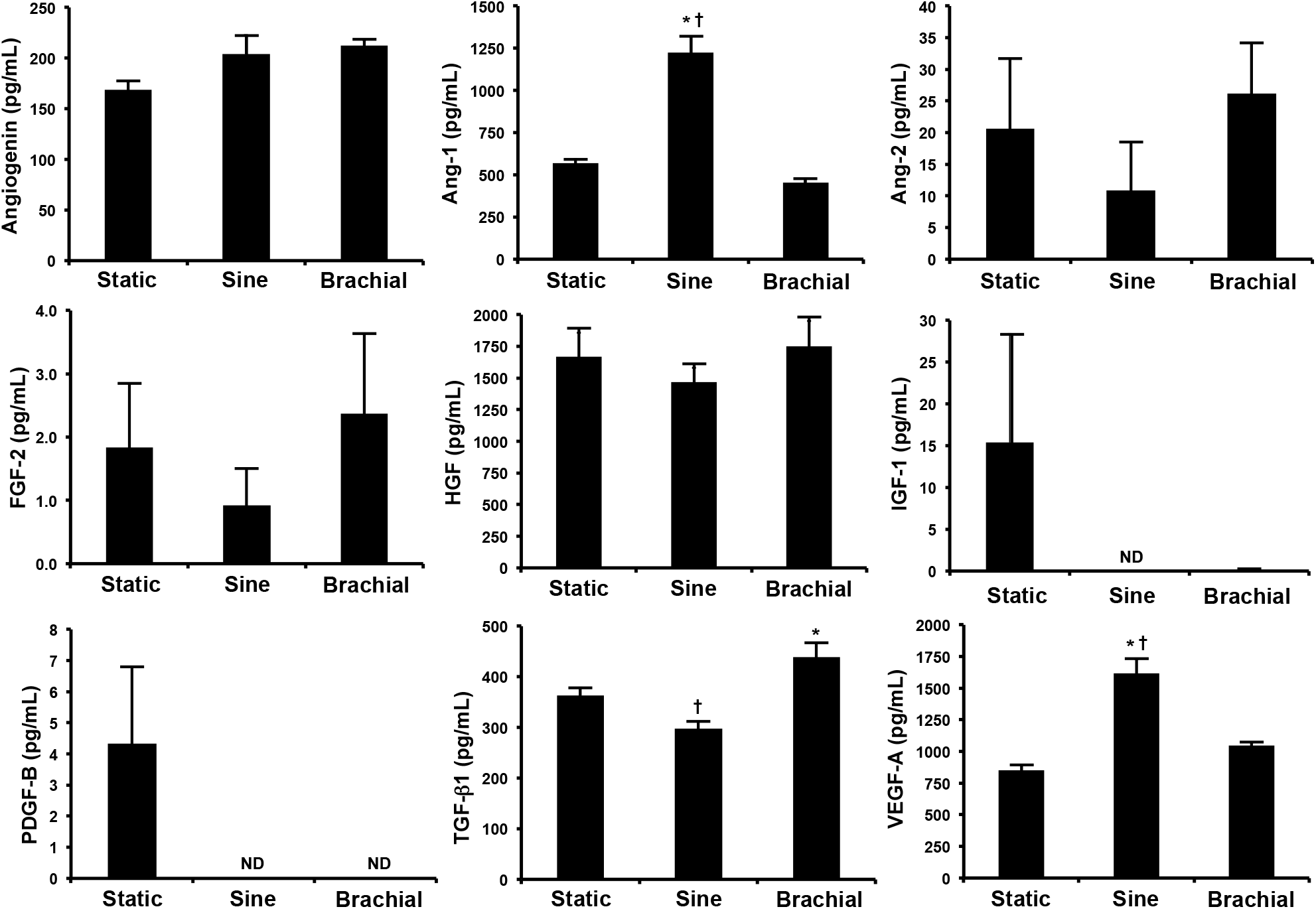
Analysis of conditioned media from MSCs treated with mechanical load (7.5% maximal strain) for 24 hours and then assayed using ELISA. \**p* < 0.05 versus static control group. †*p* < 0.05 versus brachial waveform group. ND = not detected.

**Supplemental Figure 11.**
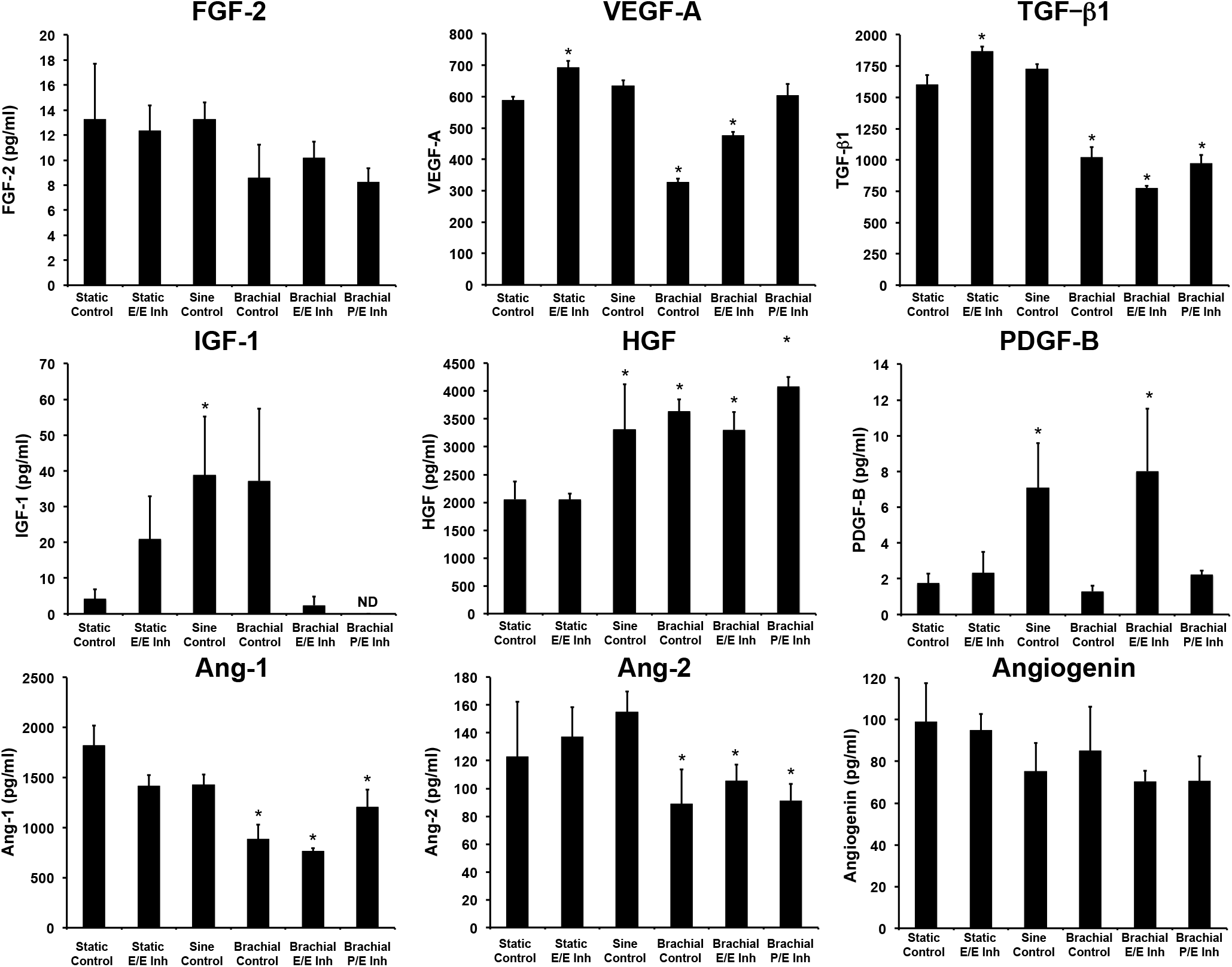
Analysis of conditioned media from MSCs treated with mechanical load (7.5% maximal strain) and pharmacological treatments for seven days and then assayed using ELISA. *\*p* < 0.05 versus static control group. ND = not detected.

**Supplemental Figure 12.**
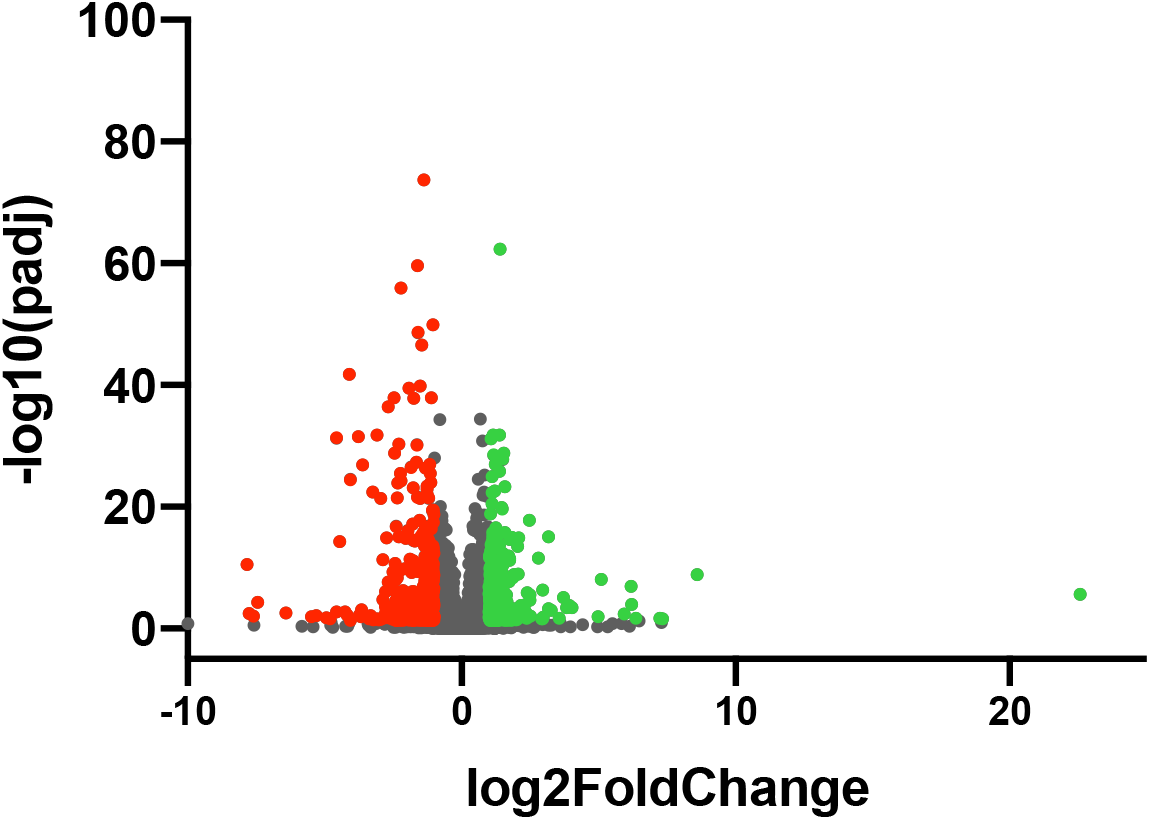
Full volcano plot for differential gene expression of the static EE group versus the static group.

**Supplemental Figure 13.**
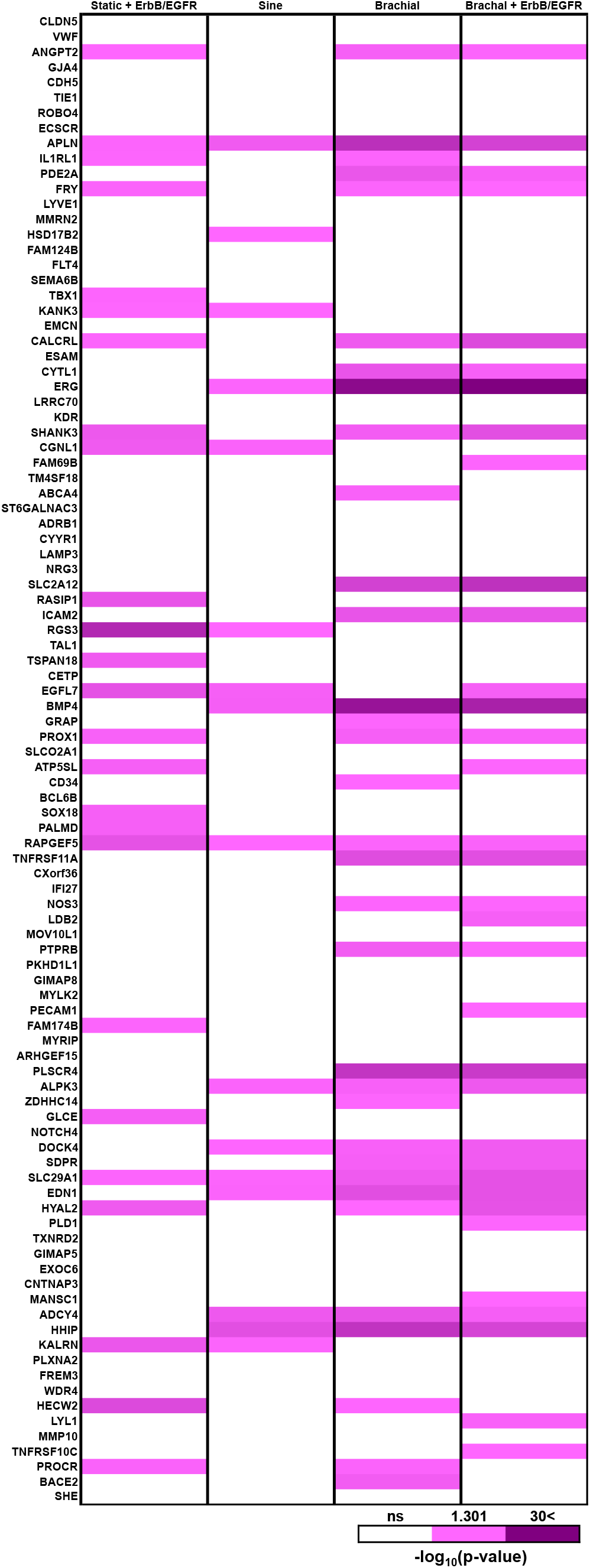
Statistical significance for the differential gene expression of endothelial phenotype related genes in comparison to the static control group.

**Supplemental Figure 14.**
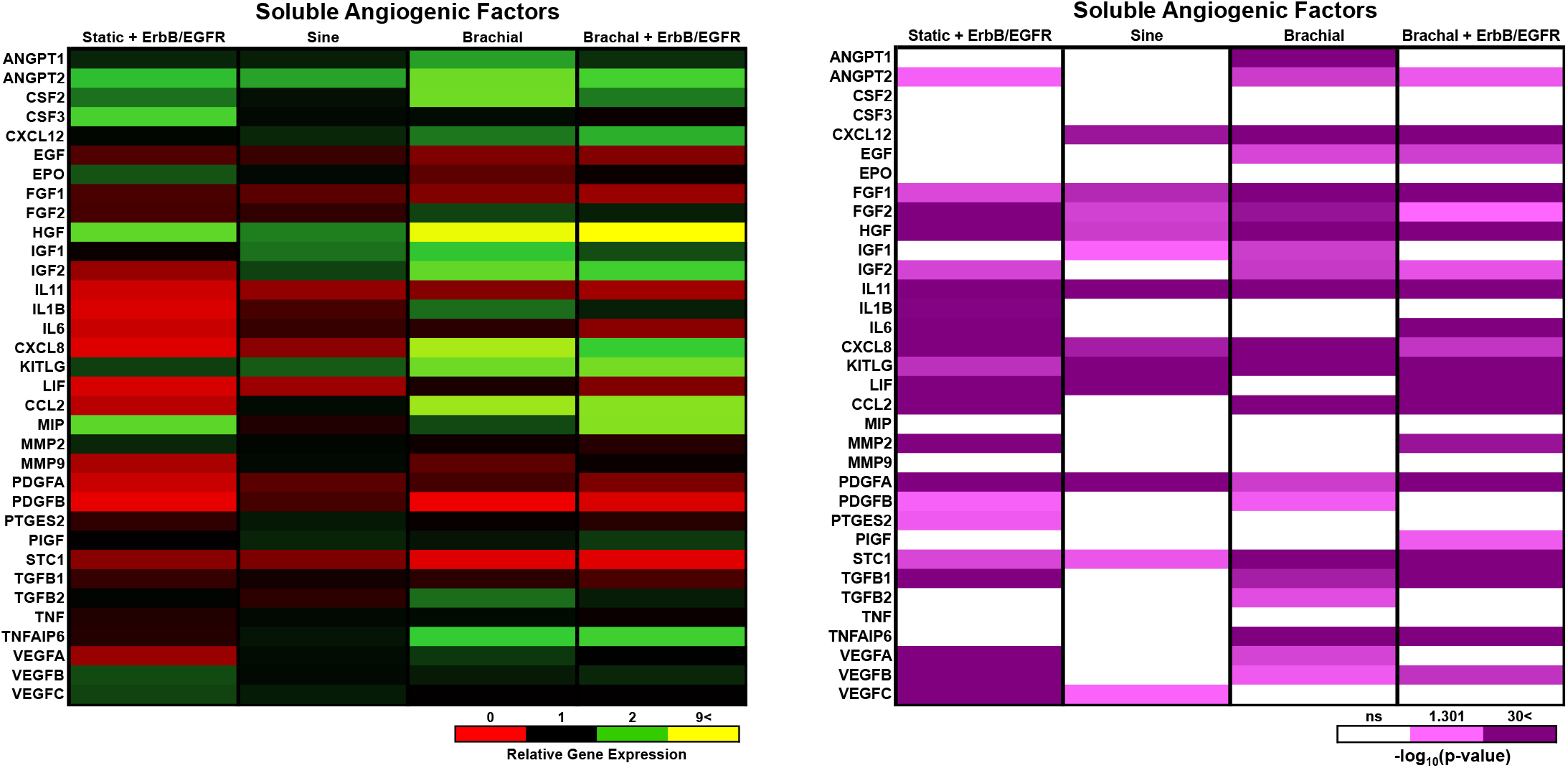
Differential gene expression for soluble angiogenic factors in comparison to the static control group.

**Supplemental Figure 15.**
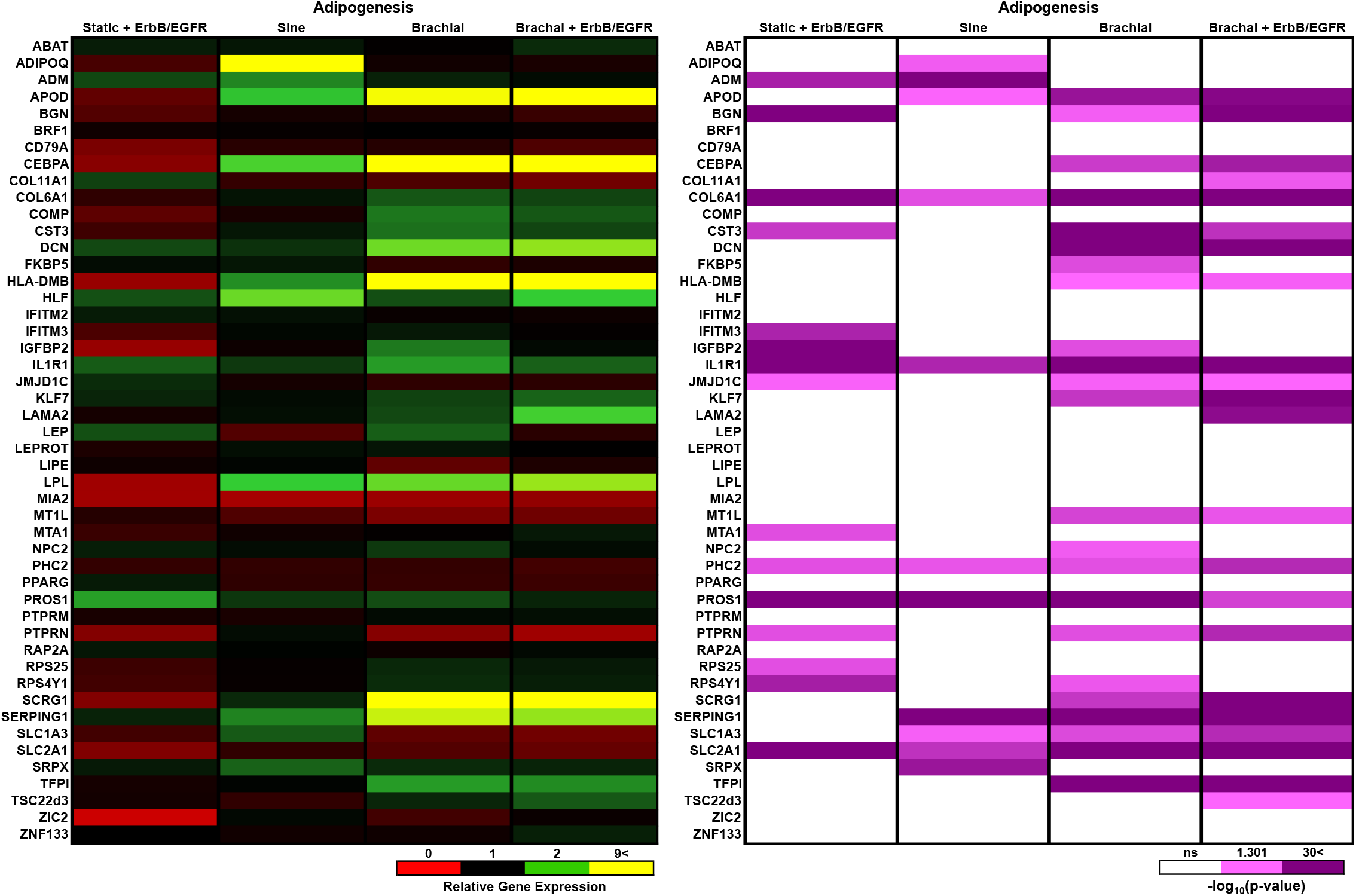
Differential gene expression for adipogenesis related genes in comparison to the static control group.

**Supplemental Figure 16.**
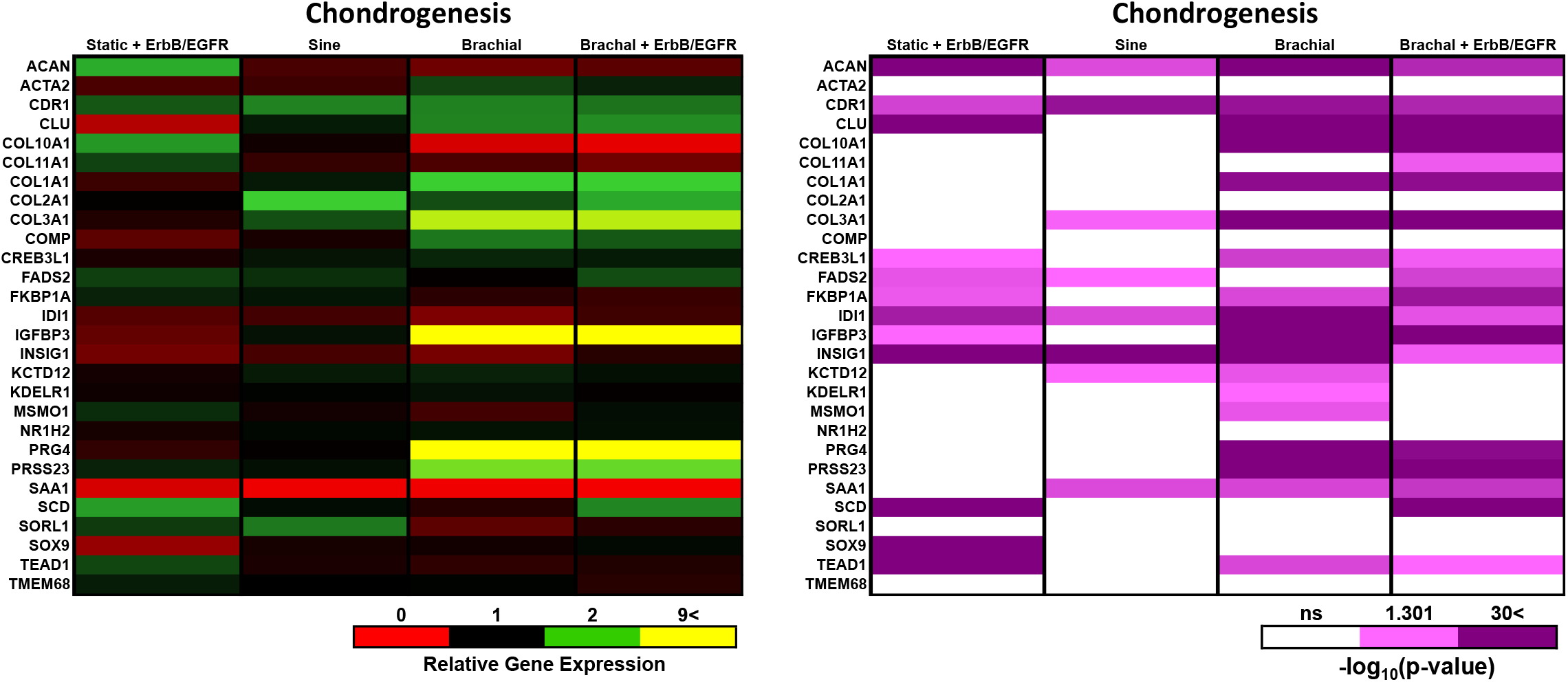
Differential gene expression for chondrogenesis related genes in comparison to the static control group.

**Supplemental Figure 17.**
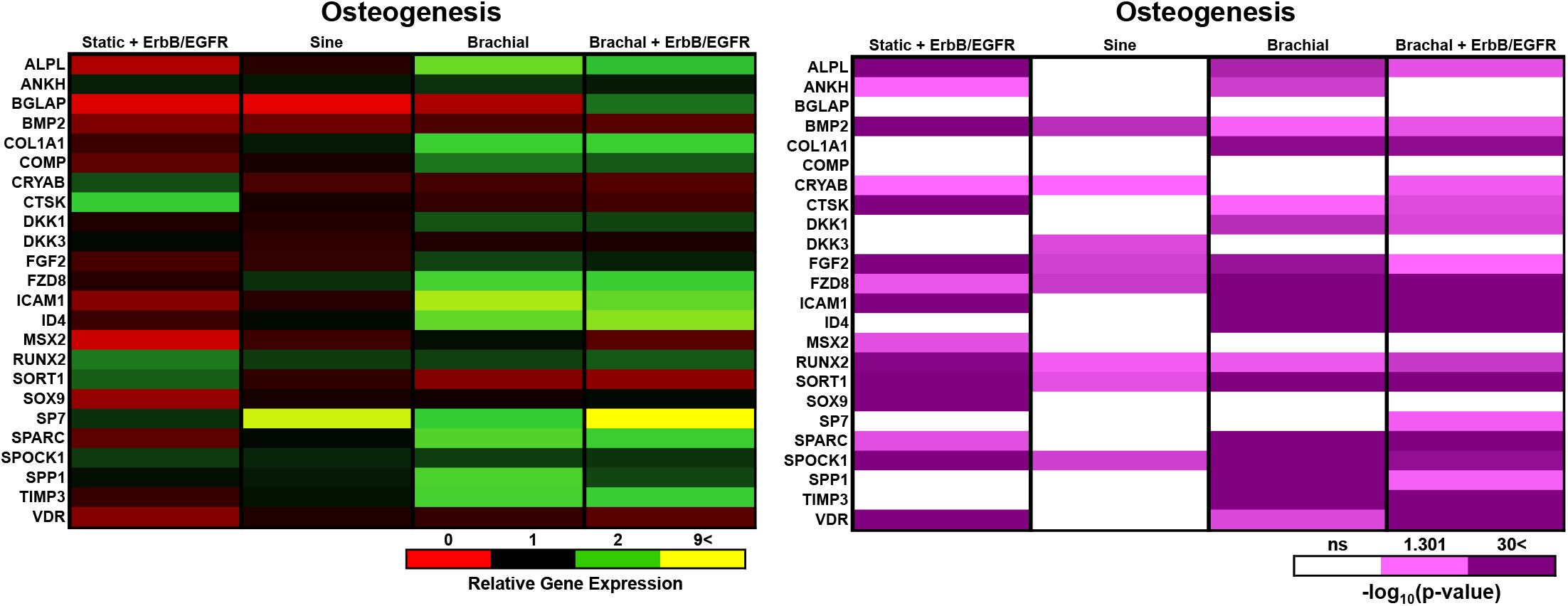
Differential gene expression for osteogenesis related genes in comparison to the static control group.

**Supplemental Figure 18.**
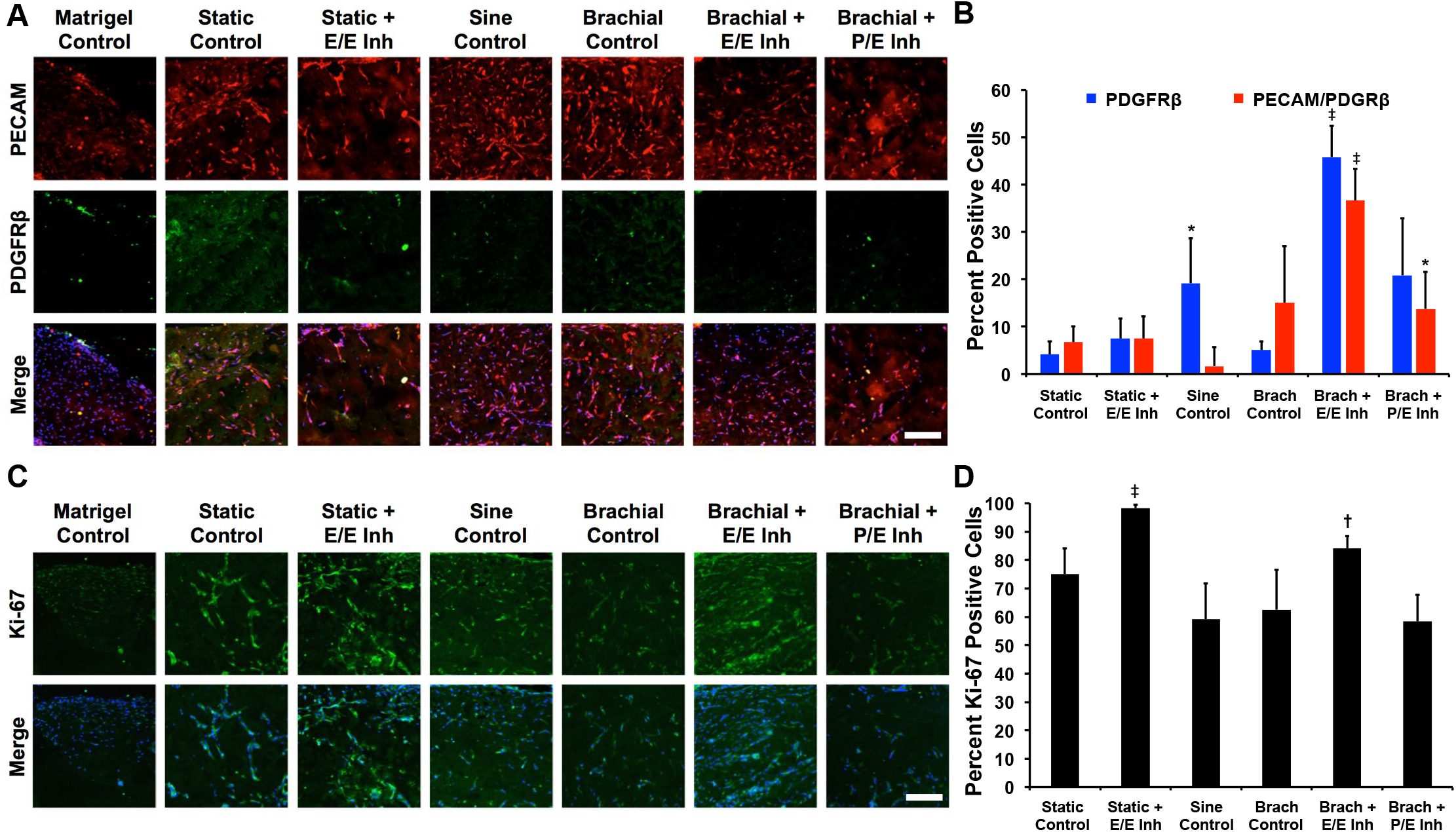
Immunostaining on histological sections the Matrigel implanted subcutaneously in nu/nu mice for 14 days. (A, B) Immunostaining and quantification for PECAM and PDGFRβ. (C, D) Immunostaining and quantification for Ki-67. *\*p* < 0.05 versus static control group. †*p* < 0.05 versus static with ErbB/EGFR inhibitor group, ‡*p* < 0.05 versus static control, static with ErbB/EGFR inhibitor, sine and brachial groups.

**Supplemental Figure 19.**
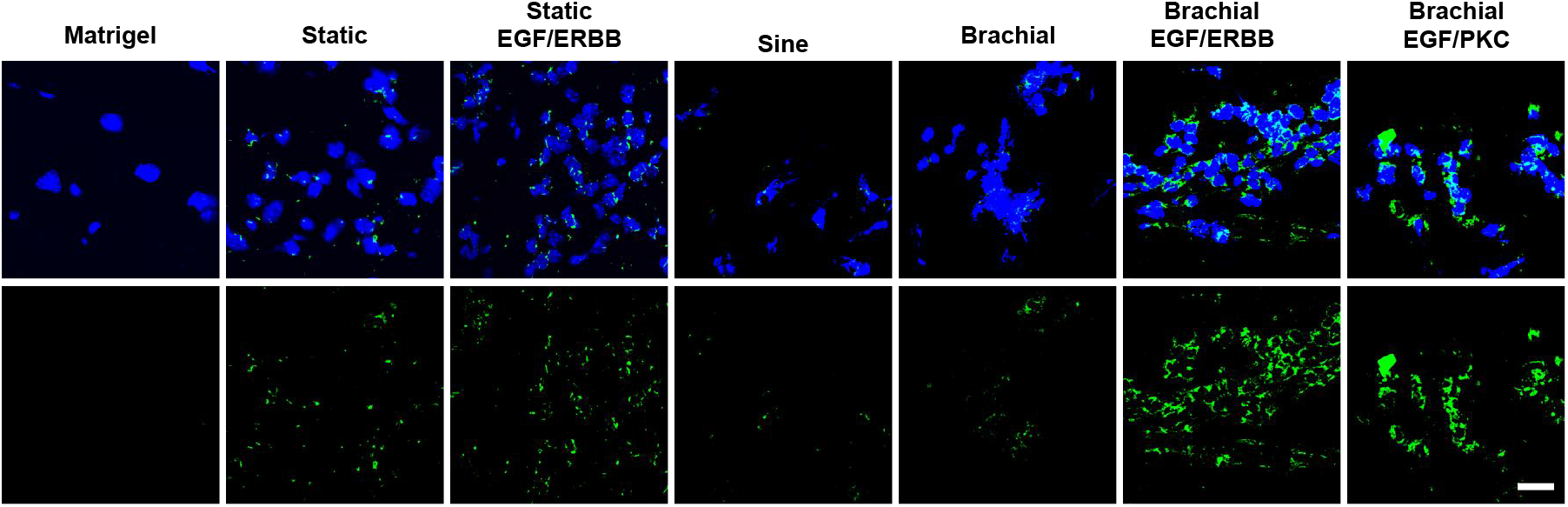
Fluorescence in situ hybridization for the human x chromosome on histological sections from mice with subcutaneous implantation of MSCs. Size bar = 200 μm.

**Supplemental Figure 20.**
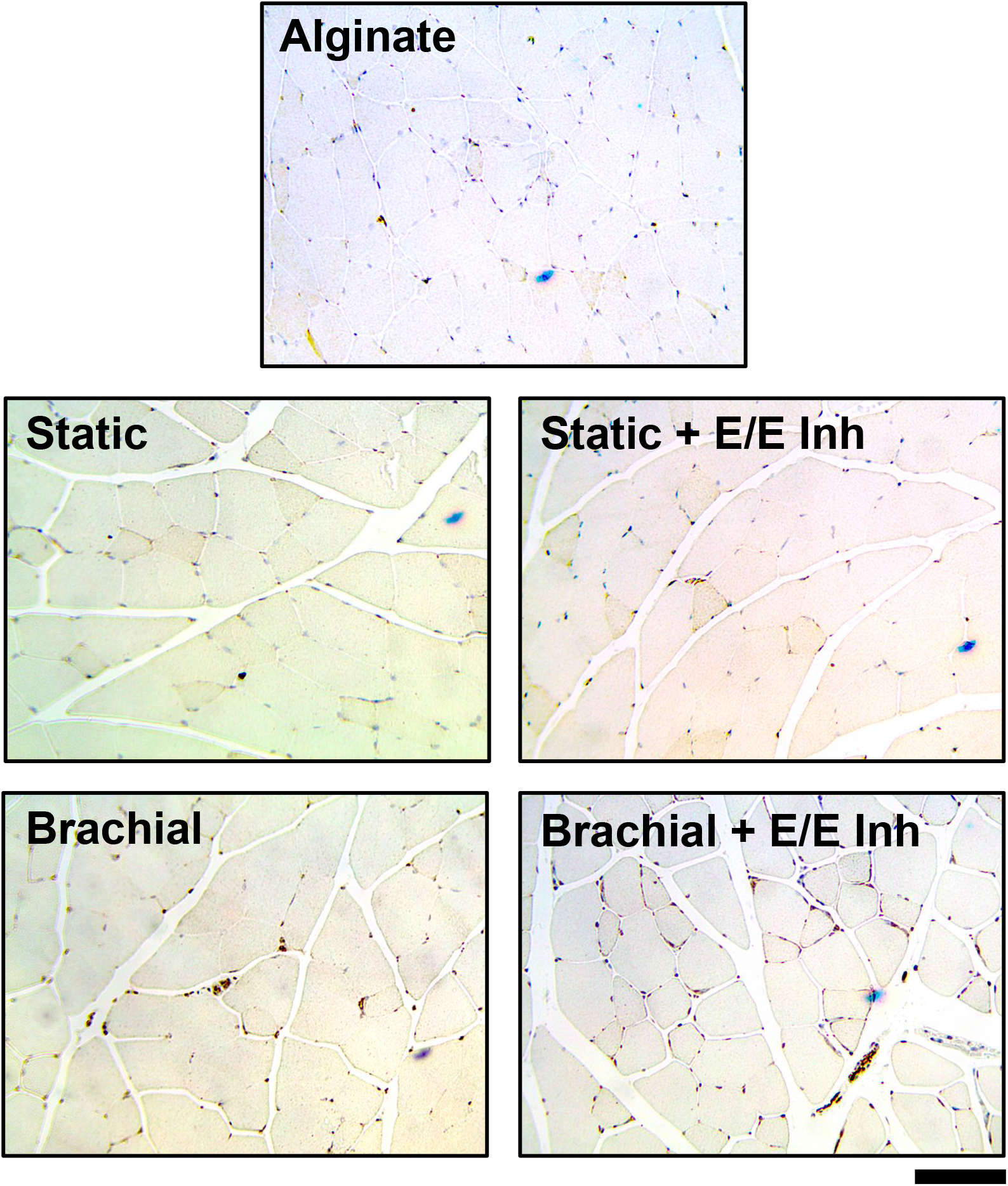
Representative images from the immunostaining analysis for PECAM-1 in the thigh muscle of the ischemic limb of mice with femoral ligation and implantation of MSCs. Size bar = 100 μm.

**Supplemental Figure 19**. Fluorescence in situ hybridization for the human x chromosome on histological sections from mice with hind limb ischemia and implantation of MSCs. Size bar = 200 μm.

**Supplemental Figure 21.**
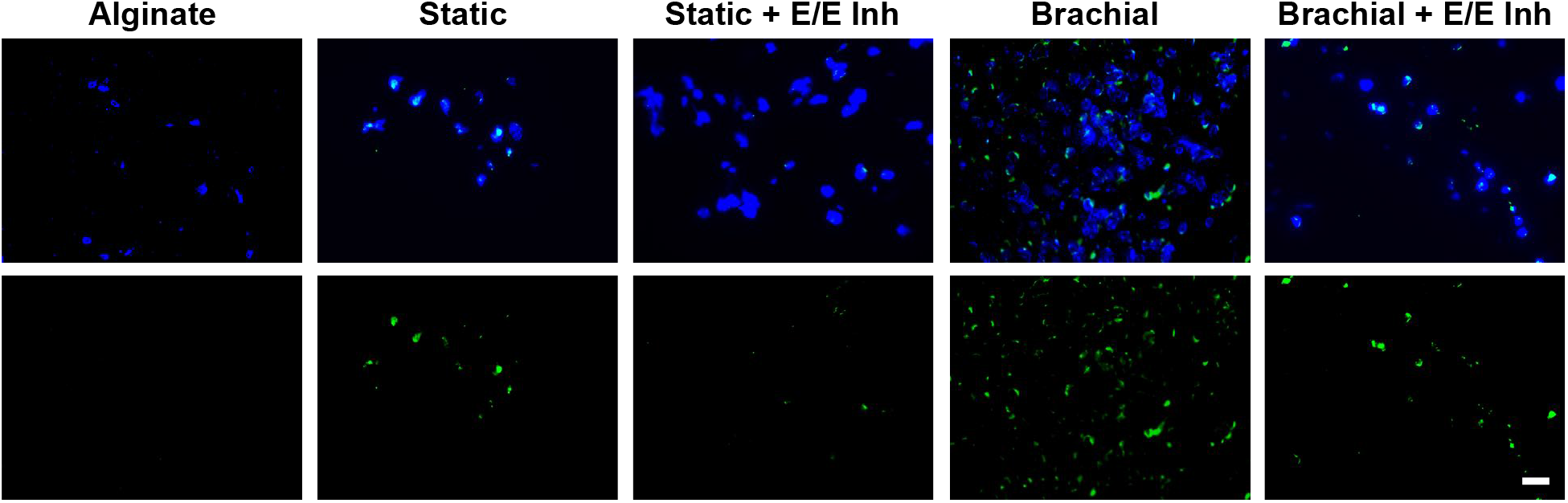
Representative images from the immunostaining analysis for PECAM-1 in the thigh muscle of the ischemic limb of mice with femoral ligation and implantation of MSCs. Size bar = 100 μm.

**Supplemental Table 1.** *Primary Antibodies/Reagents Used for Immunostaining*

| **Target Protein** | **Company** | **Catalog #** | **Species/Isotype** | **Dilution Ratio** |
| --- | --- | --- | --- | --- |
| AF 594 Phalloidin | Invitrogen | A12381 | N/A | 1:250 |
| Paxillin | Santa Cruz Biotechnology | sc-5574 | anti-rabbit | 1:50 |
| PECAM-1 | Cell Signaling | 3528S | anti-mouse | 1:100 |
| α-SMA | Abcam | ab21027 | anti-rabbit | 1:100 |
| ILK | BD Biosciences | 611803 | anti-mouse | 1:100 |
| Yap/Taz | Cell Signaling | 8418S | anti-rabbit | 1:100 |
| 14-3-3ε | Santa Cruz Biotechnology | sc393177 | anti-mouse | 1:50 |
| Smad 2/3 | BD Biosciences | 610842 | anti-mouse | 1:100 |
| Phospho-Smad2/3 | Cell Signaling | 8828S | anti-rabbit | 1:100 |
| vWF | Santa Cruz Biotechnology | 365712 | anti-mouse | 1:50 |
| Myosin IIb | Cell Signaling | 3404 | anti-rabbit | 1:100 |
| Sca-1 | Santa Cruz Biotechnology | sc-365343 | anti-mouse | 1:50 |
| Oct-4 | Cell Signaling | 2750 | anti-rabbit | 1:100 |

**Supplemental Table 2.** *Primary Antibodies Used for Immunoblotting*

| **Target Protein** | **Company** | **Catalog #** | **Species/Isotype** | **Dilution Ratio** |
| --- | --- | --- | --- | --- |
| GAPDH | Cell Signaling | 2118S | anti-rabbit | 1:500 |
| PECAM-1 | Cell Signaling | 3528S | anti-mouse | 1:500 |
| α-SMA | Abcam | ab21027 | anti-rabbit | 1:500 |
| Smad 2/3 | BD Biosciences | 610842 | anti-mouse | 1:500 |
| Phospho-Smad 2/3 | Cell Signaling | 8828S | anti-rabbit | 1:500 |
| Phospho-β Catenin | Abcam | ab27798 | anti-rabbit | 1:500 |

**Supplemental Table 3.**
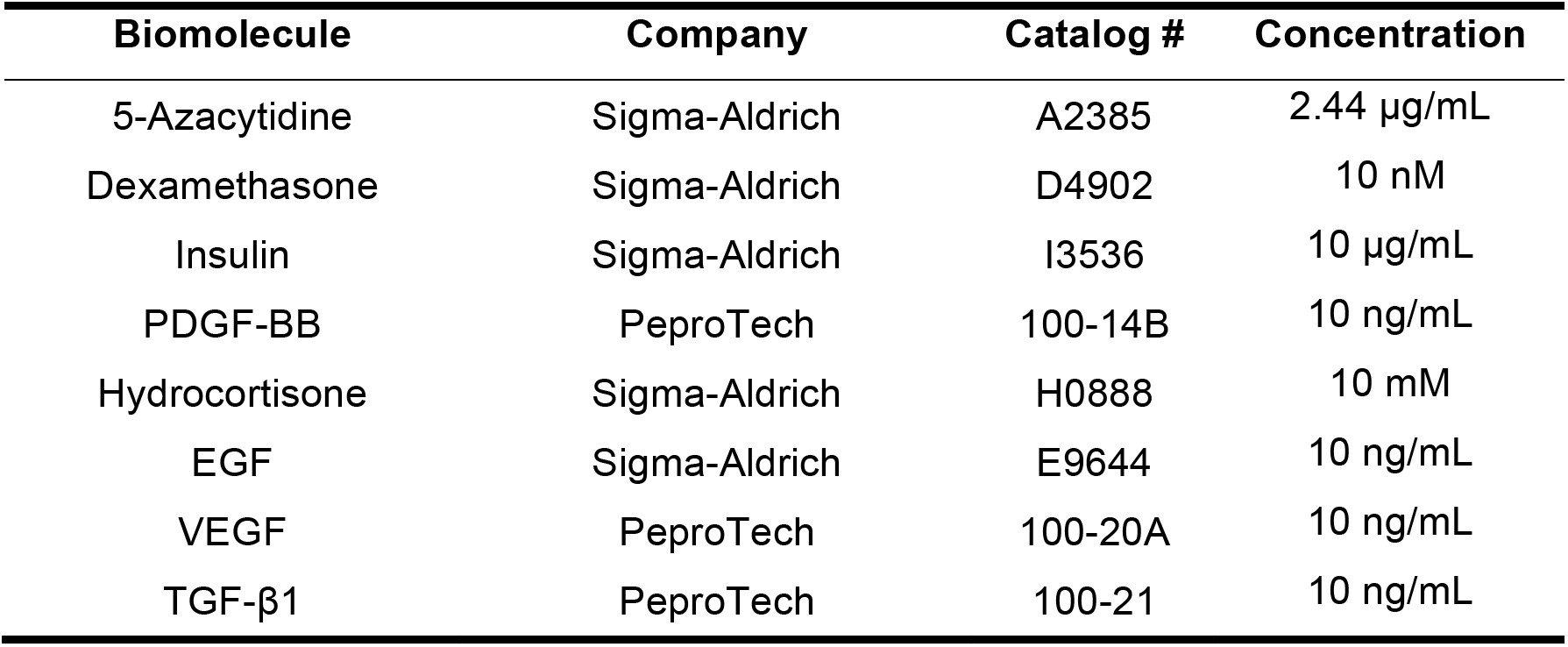
*Reagents for Treating Cells During Long Term Loading*

| **Biomolecule** | **Company** | **Catalog #** | **Concentration** |
| --- | --- | --- | --- |
| 5-Azacytidine | Sigma-Aldrich | A2385 | 2.44 µg/mL |
| Dexamethasone | Sigma-Aldrich | D4902 | 10 nM |
| Insulin | Sigma-Aldrich | I3536 | 10 µg/mL |
| PDGF-BB | PeproTech | 100-14B | 10 ng/mL |
| Hydrocortisone | Sigma-Aldrich | H0888 | 10 mM |
| EGF | Sigma-Aldrich | E9644 | 10 ng/mL |
| VEGF | PeproTech | 100-20A | 10 ng/mL |
| TGF-β1 | PeproTech | 100-21 | 10 ng/mL |

**Supplemental Table 4.** *List of Primary Antibodies Used for Flow Cytometry*

| **Target Protein** | **Company** | **Catalog #** | **Fluor. Dye** |
| --- | --- | --- | --- |
| PECAM-1 | BD Biosciences | 562855 | BV605 |
| VE-Cadherin | BD Biosciences | 565671 | BV421 |
| CD146 | BD Biosciences | 564644 | FITC |
| PDGFR-β | BD Biosciences | 558821 | PE |
| CD105 | BD Biosciences | 563466 | BV650 |
| Nestin | BD Biosciences | 561231 | PerCP/Cy5.5 |

**Supplemental Table 5.**
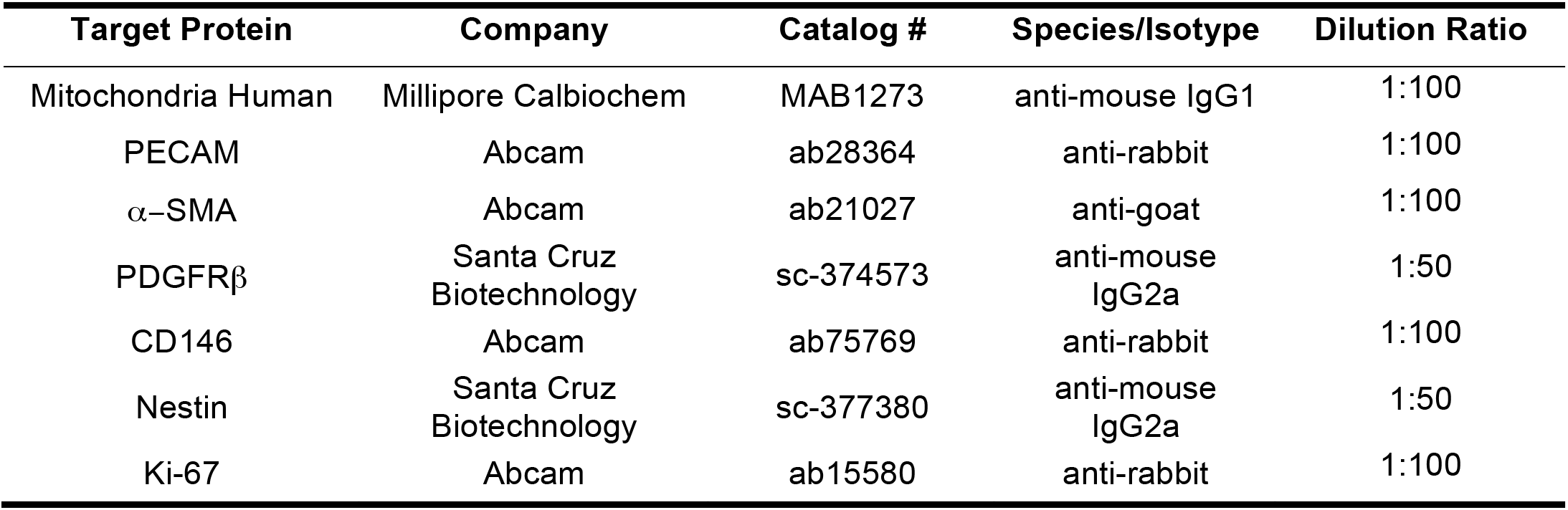
*List of Primary Antibodies Used for Tissue Immunostaining*

| **Target Protein** | **Company** | **Catalog #** | **Species/Isotype** | **Dilution Ratio** |
| --- | --- | --- | --- | --- |
| Mitochondria Human | Millipore Calbiochem | MAB1273 | anti-mouse IgG1 | 1:100 |
| PECAM | Abcam | ab28364 | anti-rabbit | 1:100 |
| α-SMA | Abcam | ab21027 | anti-goat | 1:100 |
| PDGFRβ | Santa Cruz<br>Biotechnology | sc-374573 | anti-mouse<br>IgG2a | 1:50 |
| CD146 | Abcam | ab75769 | anti-rabbit | 1:100 |
| Nestin | Santa Cruz<br>Biotechnology | sc-377380 | anti-mouse<br>IgG2a | 1:50 |
| Ki-67 | Abcam | ab15580 | anti-rabbit | 1:100 |

